# Environmental DNA in a Global Biodiversity Hotspot: Lessons from Coral Reef Fish Diversity Across the Indonesian Archipelago

**DOI:** 10.1101/2021.02.19.432056

**Authors:** Onny N. Marwayana, Zachary Gold, Paul H. Barber

## Abstract

Indonesia is the heart of the Coral Triangle, the world’s most diverse marine ecosystem. Preserving the biological and economic value of this marine biodiversity requires efficient and economical ecosystem monitoring, yet our understanding of marine biodiversity in this region remains limited. This study uses environmental DNA (eDNA) to survey fish communities across a pronounced biodiversity gradient in Indonesia. A total of 12,939,690 sequence reads of MiFish *12S* rRNA from 39 sites spanning 7 regions of Indonesia revealed 4,146 Amplified Sequence Variants (ASVs). Regional patterns of fish diversity based on eDNA broadly conformed to expectations based on traditional biodiversity survey methods, with the highest fish biodiversity in Raja Ampat and generally lower diversity in Western Indonesia. However, eDNA performed relatively poorly compared to visual survey methods in site-by-site comparisons, both in terms of total number of taxa recovered and ability to assign species names to ASVs. This result stands in a stark contrast to eDNA studies of temperate and tropical ecosystems with lower diversity. Analyses show that while sequencing depth was sufficient to capture all fish diversity within individual seawater samples, variation among samples from individual localities was high, and sampling effort was insufficient to capture all fish diversity at a given sampling site. Interestingly, mean ASVs recovered per one-liter seawater was surprisingly similar across sites, despite substantial differences in total diversity, suggesting a limit to total ASVs (~200) per one-liter eDNA sample. Combined, results highlight two major challenges of eDNA in highly diverse ecosystems such as the Coral Triangle. First, reference databases are incomplete and insufficient for effective ASV taxonomic assignment. Second, eDNA sampling design developed from lower diversity temperate marine ecosystems are inadequate to fully capture diversity of biodiversity hotspots like the Coral Triangle.

## Introduction

Indonesia is the heart of the Coral Triangle, a region in Southeast Asia that includes the Philippines, Malaysia, Timor L’Este, Papua New Guinea and the Solomon Islands. Defined by the presence of more than 500 species of scleractinian coral (Veron et al., 2009), the Coral Triangle is the world’s most biologically diverse marine ecosystem (Allen, 2008; Roberts et al., 2002; Veron et al., 2009). This remarkable diversity has made the Coral Triangle the focus of numerous biogeographic studies addressing the origins of this biodiversity hotspot (Ekman, 1953; Ladd, 1960; Woodland, 1983), as well as phylogenetic/phylogeographic studies examining speciation processes in this region (e.g., Barber & Bellwood, 2005; Barber et al., 2011; Carpenter et al., 2010; DeBoer et al., 2014; Timm, Figiel, & Kochziuz, 2008).

The biological importance of the Coral Triangle is matched only by its cultural and socioeconomic importance. More than 370 million people depend on the ecosystem goods and services of the Coral Triangle, 120 million of which directly benefit from coastal and offshore fisheries production, and marine tourism (Coral Triangle Initiative, 2009; Foale et al., 2013). For example, the fishing industry in Indonesia accounts for 21% of Indonesia’s agricultural economy and 3% of the national GDP in 2012 (FAO, 2018a). Across the Coral Triangle marine-based tourism is valued at nearly US$ 10 billion per year (NOAA, 2012).

Despite their biological and economic importance, 85% of coral reefs in the Coral Triangle are threatened or extremely threatened (Burke et al., 2012). Major threats include habitat degradation, marine pollution, increasing global demand for marine products, coastal deforestation, and unsustainable fishing (Hoegh-Guldberg et al., 2009). Not only are reefs degrading, but even healthier reef systems like Raja Ampat, Indonesia, larger members of the fish community, such as grouper and Napoleon wrasse, have greatly reduced population sizes (Allen, 2003) as do reef sharks (Sembiring et al., 2015). These losses don’t just impact biodiversity, they also impact ecosystem function and the human communities that depend on them (Burke et al., 2012). The latter is particularly concerning because nations like Indonesia lack sufficient terrestrial habitat to compensate for lost marine productivity and have insufficient livestock production is to meet local protein demands (Permani et al., 2016). As such, it is critical to preserve the Coral Triangle marine ecosystems and the socio-economic benefits that these ecosystems provide.

One important aspect of sustainability in the Coral Triangle is effectively monitoring change over time. However, there are two major challenges to effective monitoring. First, research effort in the Coral Triangle is not proportional to its exceptional biodiversity (Barber et al., 2014; Fisher et al., 2011; Keyse et al., 2014) leading to an incomplete understanding of marine biodiversity in this ecosystem. A second challenge is that the most common biodiversity survey method, particularly for corals and fishes, is visual census. Visual census is expensive, time intensive, potentially biased by observer skill, and can overlook rare or cryptic taxa (Edgar et al., 2004). The latter two are particularly problematic in Indonesia because areas like Raja Ampat have at least 1,704 species of marine fish (Allen, 2002), and few scientists have the taxonomic expertise to conduct such surveys. Furthermore, logistical issues and methodological biases hinder researchers’ ability to do time-series visual surveys, limiting our ability to understand how these ecosystems are changing over time, precluding the collection of monitoring data that is critical to inform marine conservation efforts. As such, it is essential to develop novel methods that are 1) efficient, 2) inexpensive, 3) require no specific taxonomic expertise, and 4) are amenable to large scale spatial and temporal sampling.

Environmental DNA (eDNA) is a rapidly evolving approach to the study of biodiversity (Deiner et al., 2017). eDNA is widely applied to document the presence of endangered species (e.g., crested newt; Biggs et al., 2015; Rees et al., 2014, 2017), invasive species (e.g., the American bullfrog; Dejean et al., 2012; Rees et al., 2014; Takahara et al., 2013) and diversity of freshwater fish communities (Rees et al., 2014; Thomsen et al., 2012b). eDNA also shows great promise for studies of marine biodiversity (Closek et al., 2019; Cordier et al., 2017), recovering commercially valuable species as well as species rarely or never recorded by conventional monitoring methods (Thomsen et al., 2012a). Port et al. (2016) showed that eDNA can differentiate fish communities between adjacent habitats within a coastal kelp forest ecosystem even in the presence of strong water movement(Jeunen et al., 2019), and more recently, Gold et al (2021) showed that eDNA can differentiate fish communities inside and outside of marine protected areas. Importantly, a common theme of marine eDNA studies is that eDNA recovers a greater diversity of taxa than traditional visual surveys (e.g., Gold et al., 2020; Kelly et al., 2014, 2017), making it a particularly attractive approach for biodiversity monitoring in regions like the Coral Triangle (Juhel et al., 2020).

While eDNA may revolutionize monitoring of marine biodiversity, to date most marine eDNA studies have focused on relatively low diversity temperate marine environments (Bohmann et al., 2014; McElroy et al., 2020; Rees et al., 2017; Thomsen et al., 2012a). Similarly, most coral reef eDNA studies to date have occurred in Japan, Central Pacific, Western Australia, and the Red Sea (Alexander et al. 2020;Bessey et al. 2020; DiBattista et al. 2017; Lafferty et al. 2020; Oka et al. 2020;Stat et al 2017; Stat et al. 2019; West et al. 2020), where fish biodiversity is far lower than the Coral Triangle (Bellwood & Meyer, 2009; Roberts et al., 2002). Most recently, Juhel et al. (2020) used eDNA to examine fish diversity in Raja Ampat, Indonesia, home to the world’s most diverse reef fish fauna (Allen, 2002), reporting a strong correlation between eDNA results and checklists from visual data. However, species accumulation curves indicated that eDNA greatly under sampled species diversity, suggesting that the methods, sampling design, and bioinformatics approaches employed in temperate eDNA studies may not be equally effective in megadiverse tropical regions like the Coral Triangle.

This study employs eDNA to examine the distribution and diversity of marine fishes across the Indonesian archipelago and compare these results to patterns of region diversity resulting from conventional visual census that show a strong biodiversity gradient. Specifically, we test the potential of eDNA to provide a simple, cost-effective method for assessing and monitoring fish communities in a threatened global biodiversity hotspot by asking (1) whether eDNA effectively captures regional patterns of fish biodiversity across the Indonesian Archipelago, and (2) whether eDNA captures more fish diversity than traditional survey methods, including rare and cryptic taxa, as is common in lower biodiversity marine ecosystems.

## METHODS

### Sampling Sites

We collected eDNA samples across Indonesia, spanning a strong fish biodiversity gradient (Bellwood & Meyer, 2009; Roberts et al., 2002). Sampling focused on three regions (Figure 1), including: 1) outside the Coral Triangle in Western Indonesia (Aceh and Batam-Bintan), 2) lower diversity regions of the Coral Triangle in Central Indonesia (Derawan and Wakatobi), and 3) high diversity regions of the Coral Triangle (Eastern Indonesia: Lembeh Strait, Ternate, and Raja Ampat) that have the world’s highest reef fish biodiversity (Allen & Werner, 2002; Bellwood & Meyer, 2009; Roberts et al., 2002).

### eDNA Sampling

To assess marine fish diversity with eDNA, we employed a hierarchical sampling design across 7 regions (Table S1). Each sample consisted of one liter of seawater that was collected on SCUBA at depths between 11-15 m, 1 m off the reef, to minimize variation in community composition associated with depth. Following standard sampling protocols used in temperate ecosystems (Miya et al., 2015), we collected three one-liter water samples at each sampling site to maximize species diversity and to account for fine-scale heterogeneity in local eDNA signatures. To further maximize species diversity, we sampled multiple sites within each region, with sampling sites separated by at least five kilometers to capture spatial variability in habitat and eDNA signatures (Andruszkiewicz et al., 2017; Miya et al., 2015), except for Wakatobi, where sites were 2.5 km distant. For example, on the island of Derawan, we took three one-liter water samples from each of four sites, totaling 12 individual eDNA samples. To compare eDNA to visual census surveys, we selected sites in Raja Ampat where Allen and Erdmann conducted intensive visual fish surveys (pers. comm.).

To isolate eDNA from seawater samples, we filtered one liter of seawater through a 0.22-micron Sterivex™ filter (Millipore®, SIGMA MILLIPORE) following the methods of Miya et al. (2015) and Curd et al. (2018) with one key modification: we collected individual water samples in sterile one liter Kangaroo™ Gravity Feeding Bags (similar to intravenous drip bags) that allow for gravity filtration through the Sterivex columns, a method ideally suited to remote field locations. In addition, we filtered one liter of bottled drinking water at each location to serve as a “field blank” negative control. Filters were stored in a −20° C freezer until eDNA was extracted at the the Indonesian Biodiversity Research Center (now “Bionesia”).

### eDNA Extraction, Amplification, and Sequencing

We extracted eDNA samples and blanks using the DNeasy Blood & Tissue Kit (QIAGEN, Hilden, Germany) following the modified extraction protocol of Spens et al. (2017), adding 720 μL of ATL buffer and 80 μL of proteinase K directly into the filter cartridge. Extracted DNA was then stored at −20°C and transported to UCLA for amplification and library preparation.

We amplified extracted eDNA using the Multiplex PCR Kit (QIAGEN, Hilden, Germany), targeting two DNA markers, the MiFish Teleost and MiFish Elasmobranch, *12S* rRNA mitochondrial DNA (mtDNA) primers (Miya et al., 2015). We conducted each individual eDNA sample via Polymerase Chain Reactions (PCR) in triplicate to account for potential PCR bias (Andruszkiewicz et al., 2017; Miya et al., 2015; Taberlet et al., 2012). Each PCR reaction consisted of 12.5 μL Qiagen 2x Master Mix, 2.5 μL (2 mM) of the primer, 6.5μL nuclease free water, and one μL the DNA extract. Thermocycling parameters utilized a touchdown protocol, beginning with a 15-minute pre-denaturation step at a 95 °C, followed by a touchdown thermocycling profile consisting of 30 seconds denaturing at 94 °C, 30 seconds annealing at 69.5 °C, and 30 seconds extension at 72 °C, with the annealing temperature dropping by 1.5 °C per cycle until 50 °C. Following this initial touchdown phase, the main cycle consisted of 25 cycles of 94 °C for 30 seconds for denaturation, 50 °C for 30 seconds for annealing and 72 °C for 45 seconds for extension, concluding with a 10-minute final extension at 72 °C. We confirmed successful PCR reactions through electrophoresis of 5μL products for 30 minutes at 150 volts on 2% agarose gels stained with 6x SYBR™ Green (ThermoFisher Scientific, Waltham, MA, USA) for visualization.

To create the sequencing libraries, we pooled the triplicate PCR products from each individual eDNA sample into a single tube and purified these pooled PCR products using the serapure bead protocol (Faircloth et al. 2014). Next, we quantified the DNA concentration (ng/μL) of each pooled PCR sample using the Qubit Broad Range dsDNA assay (ThermoFisher, Waltham, MA, USA) following the manufacturer protocol, and then then subsequently normalized pooled PCRs to equimolar concentration. We then used the Nextera DNA Library Preparation Kit (Illumina, San Diego, CA, USA) to index each pooled sample using a unique combination of Nextera i5 and i7 primers in a second PCR reaction, following the manufacturer protocol (Curd et al., 2018). The indexing PCR reaction consisted of 12.5 μL Kapa High Fidelity Master Mix (Roche, Basel, Switzerland) 0.625 μL of 1 μM i5 Illumina Nextera indices, 0.625 μL of 1 μM i7 Illumina Nextera indices, and 11.25 μL of PCR product for a total of 10ng of DNA. The indexing PCR thermocycling parameters began with an initial denaturation of 95 °C for 5 minutes, followed by 8 cycles of: 98 °C denaturation for 30 seconds, 56 °C annealing for 30 seconds, and 72 °C extension for 3 minutes, ending with a 72 °C extension for five minutes. To ensure the indexing PCR was successful, we electrophoresed indexed PCR products at 120 V for 45 minutes on a 2% agarose gel. We then created the final sequencing library by combining the cleaned and quantified indexed PCR products in equal concentration of 10 ng/μl per sample. Libraries were pooled by barcode resulting in two pooled libraries. We also note that the final MiFish Elasmobranch library contained samples collected from surf zones in Southern California which were removed at the end of the decontamination process described below. The final libraries were sequenced at the UC Berkeley sequencing core on an Illumina MiSeq platform utilizing 300 base pair paired end sequencing.

### Bioinformatics and Data Analysis

To determine fish community composition, we used the *Anacapa Toolkit* (version: 1) to conduct quality control, amplicon sequence variant (ASV) parsing, and taxonomic assignment using user-generated custom reference databases (Curd et al., 2018). The *Anacapa Toolkit* sequence QC and ASV parsing module relies on *cutadapt* (version: 1.16) (Dobin et al., 2013), *FastX-toolkit* (version: 0.0.13) (Gordon & Hannon, 2010), and *DADA2* (version 1.6) (Callahan et al., 2016) as dependencies, and the *Anacapa classifier* modules relies on *Bowtie2* (version 2.3.5) (Langmead & Salzberg, 2012) and a modified version of BLCA (Gao, et al., 2017) as dependencies. We processed sequences using the default parameters and assigned taxonomy using a *CRUX*-generated reference database supplemented with additional *12S* reference sequences generated by the Smithsonian (GenBank Accessions Table S2). We note that *CRUX* relies on *ecoPCR* (version: 1.0.1) (Boyer et al., 2016), *blastn* (version: 2.6.0) (Altschul et al., 1990; Camacho et al., 2009), and *Entrez-qiime* (version: 2.0) (Baker, 2016) as dependencies.

Raw ASV community table was decontaminated following Kelly et al. (2018) and McKnight et al. (2019) (See Supplemental Methods). However, we did not conduct site occupancy modeling as we were specifically interested in exploring under sampling of total ASV diversity. We then transformed all read counts into an eDNA index for beta-diversity statistics (Kelly et al., 2019). Because of noted issues with incomplete reference databases (Juhel et al., 2020), analyses focused on total ASVs instead of ASVs successfully assigned to species.

### Patterns of Biodiversity Across Indonesia

We conducted analyses of fish biodiversity in a hierarchical fashion. First, we examined diversity at the level of individual sample replicate, where fish diversity was represented by eDNA sequences amplified from each of the three one-liter water samples at an individual sampling site. Second, we examined diversity at the site level, where fish diversity was represented by combining replicate eDNA sequences amplified from each site (Table S1). Lastly, we examined diversity in each of the seven sampled regions by combining diversity across all sites sampled. To compare ASV recovery in lower diversity temperate marine ecosystems to high diversity reefs of the Coral Triangle, we compared the above to eDNA fish diversity data from Scorpion Bay, Santa Cruz Island, California (Gold et al., in press) as samples were collected, processed, an analyzed using the same methods outlined above. All analyses were conducted in in *R* (R Core Team 2020).

We examined patterns of alpha diversity by exploring species richness at each hierarchical level. We then compared sample coverage estimates by conducting ASV accumulation curves using the iNext package (version: 2.0.20) in *R* (Hsieh et al., 2016). Specifically, we explored the accumulation of ASVs 1) across all replicate samples within each site, 2) within all replicate samples within each region, and 3) within all replicate’s sites within each region. We then explored betadiversity patterns across sample replicates, sites, and regions by calculating jaccard-binary dissimilarity indices and running PERMANOVA and companion homogeneity of dispersion test (*adonis* and *betadisper* functions from *vegan* (version: 2.5-6) in *R*). Results of the PERMANOVA were visualized using a nonmetric multidimensional scaling (NMDS) through the *phyloseq* (version: 1.32.0) and *vegan* packages in R (R Core Team, 2020).

To explore the scale of species turnover across the biogeographic regions, we conducted zeta-diversity analyses using the *zetadiv* (version 1.2.0) package (Latombe et al., 2017). Zeta diversity measures the degree of overlap between any set of observed communities (Hui & McGeoch, 2014), in contrast to betadiversity which only compares pairwise overlap (Hui et al., 2014; Latombe et al., 2017; McGeoch et al., 2019). Importantly, the zeta diversity framework allows for the calculation of diversity metrics across multiple samples and sampling strata to better understand patterns of biodiversity (McGeoch et al., 2019), particularly the scale and shape of community turnover (Hui et al., 2014; McGeoch et al., 2019; Simon et al., 2019). From this framework there are two important zeta diversity metrics: zeta diversity decay and species retention rates (McGeoch et al., 2019). Zeta diversity decay measures the decreasing number of shared taxa across greater observed communities (McGeoch et al., 2019). For small numbers of samples (n<5) this can be visually represented as the center of the Venn Diagram between multiple observed communities (i.e., individual eDNA samples).

We compared species retention rates and zeta diversity decay (i.e., the decreasing number of shared taxa across more observed communities) across 1) all replicate samples within each site, 2) within all replicate samples within each region, and 3) within all replicate sites within each region. In particular, we compare the rate of zeta diversity decay and species retention rates across the region to identify sites with lower community turnover across sampling strata. We further explore zeta diversity decay over distance to explore the scale of community turnover across sample replicates, sites, and regions through eDNA data.

### Comparison of eDNA Survey and Visual Census

To compare the efficacy of eDNA as a biodiversity survey tool relative to traditional visual census surveys in high diversity marine ecosystems,, we compared eDNA metabarcoding results to high-intensity visual census data generated by Gerry Allen and Mark Erdmann (pers. comm.), in Raja Ampat. These surveys involve multiple divers spending 6 hours per day over 10-21 and have the ultimate goal of surveying as much of the fish community as possible. Because the performance of eDNA can be hindered by incomplete reference databases, we compared the effectiveness of eDNA and visual census to both total ASVs as well as ASVs identified to species.

## RESULTS

### Raw eDNA Sequences

From 119 eDNA samples collected between June-August 2017, we generated a total of 15,219,431 raw sequence reads. After processing via the Anacapa Toolkit and subsequent decontamination, these sequences yielded a total of 4,688,144 MiFish Elasmobranch reads representing 1,783 ASVs and 8,251,546 MiFish Teleost reads representing 2,363 ASVs. Sample read depth ranged from 11,178 to 308,459 reads (mean 108,737 reads) and ASVs per sample appeared to saturate for all samples (Figures S1 & S2, Table 1). Of the total 4,146 ASVs, we were only able to successfully assign species level taxonomy to 2,309 ASVs (55.7%), genus level to 483 ASVs (11.6%), family level to 66 ASVs 1.6%), order level to 7 ASVs (0.16%), class level to 14 ASVs (0.32%), and 1,267 (30.6%) could not be assigned to any taxonomic rank (Tables S3 & S4). From these assignments we identified 642 unique species, 333 genera and 102 families of teleost and 20 unique species, 16 unique genera, and 10 families of elasmobranchs.

### Regional Pattern and Identification of Amplicon Sequence Variants

Fish diversity was highest in Eastern Indonesia, as expected, with Raja Ampat having the highest number of unique ASVs (n= 1,706), followed by Lembeh Strait (n= 1,292). However, the other Eastern Indonesia site, Ternate, had much lower numbers (n= 695), similar to Wakatobi (n= 717) and Derawan (n= 799) (Figure 2b). As expected for regions outside the Coral Triangle, Bantam Bintam had the lowest diversity (n= 196); however, ASV diversity in Aceh (n= 879) was more similar to regions inside the Coral Triangle.

Total ASV per sample varied from a low of 196 in Batam-Bintam to a high of 1,706 in Raja Ampat. Accounting for sampling effort, we found a significant difference in ASVs per sample replicate across regions (ANOVA p <0.00001; Figure 2c) with Batam Bintang having significantly fewer ASVs per replicate sample than Aceh, Raja Ampat, and Wakatobi (Tukey HSD, p <0.05; Table S5), whol;e Raja Ampat had significantly greater diversity per sample replicate than Lembeh Strait (Tukey HSD, p <0.05; Table S5). Results showed significant differences in the mean number of ASVs per sites (ANOVA and Tukey HSD, p<0.05, Figure S3, Table S6) and a significant difference in the total number of ASVs per site across regions with Batam Bintam having significantly lower diversity than Aceh, Raja Ampat, Wakatobi, and Ternate (ANOVA, p<0.05, Figure 2a, Table S7). While all sampling localities, except Batam-Bintam, had higher total diversity and per site diversity compared to Southern California, ASV diversity per sample was remarkably similar across all locations.

ASV accumulation curves across all replicate samples showed only Batam-Bintam approached saturation (95.7% sample coverage) (Figure 3a, Table S8), indicating sufficient sampling and sequencing depth. In contrast, sample coverage estimates within the other 6 regions ranged from 44.3% to 65.6% for all samples surveyed. This pattern was also true for ASV accumulation curves at the sites level (Figures S4-S10). Batam-Bintam had a site coverage estimate of 92.5% while site coverage from all other regions ranged from 28.4% to 64.0% (Figure 3b, Table S9). Likewise, ASV accumulation curves across sample replicates within each site only saturated at Batam-Bintam (90.0-92.6% sample coverage) with sample coverage at all other sites ranging from 14.6%-60.7% (Table S10). Estimates show that lower-diversity regions (e.g., Batam-Bintam) need ~66 samples to capture nearly all fish diversity present (99.9% sample coverage), while the higher-diversity regions (e.g., Raja Ampat) require >300 samples to obtain 99.9% sample coverage. In contrast to the above results, in all cases individual eDNA samples had sufficient read depth to saturate ASV recovery (Figures S1 & S2). Together, these results demonstrate substantial under-sampling of diversity at the site, and region levels, but not at the individual sample level.

Despite under-sampling, region and site explained 15.6% and 30.7% of variation in fish communities across Indonesia (PERMANOVA, p <0.001, Table S11). Results also show significant differences in dispersion of fish communities between sites and regions, largely driven by low variance of dispersion in Batam-Bintang (betadisper, p <0.001, Table S11), consistent with the results of the species accumulation curves. Visualization of these results through NMDS ordination showed substantial overlap among all seven regions, albeit with Batam-Bintam and Aceh having the most distinct clustering of eDNA samples (Figure 4).

Zeta diversity decay and species retention showed that that Batam-Bintam had significantly lower zeta diversity decay across both samples and sites than the other regions, strongly indicating greater community overlap in these Western regions (Figure 5). Zeta diversity across replicate samples decayed to near zero for all other regions at an order of 3, corresponding with the most frequent number of sample replicates taken at each site. Zeta diversity across sites similarly decayed to zero around 3 sites for all regions except Bata-Bintam and Wakatobi which displayed much higher community overlap between sites.

Comparing sample and site level zeta diversity decay across distance showed that sites within less than 10 km had the greatest degree of community overlap, decreasing to near zero beyond (Figure S11), suggesting that only sample replicates taken at the exact same location and time had low community turnover. The rapid zeta diversity decay over distance helps explain the high degree of community overlap in Wakatobi as all sites were within 2.5 km.

We observed higher species retention rates across sites in Western Indonesia, indicating lower community turnover at Aceh and Batam-Bintam than in Raja Ampat and Lembeh Strait. Species retention rates across all samples in the region found similar patterns except for Batam Bintam which had an initial increase in species retention rates. These patterns correspond to a lack of many common species between samples and sharp community turnover. However, Batam Bintam also demonstrated the greatest site species retention rates and lowest zeta decay suggesting these results are indicative of a higher degree of shared species within each site than within the region.

Comparing species retention rates within regions across all samples revealed patterns of initial increase followed by steep declines for all regions except Aceh and Wakatobi (Figures S12-18). In contrast, the saturation of species retention within Aceh and Wakatobi indicates that there are a handful of common species across all replicate samples, and that Aceh has a greater total number of shared common species than Wakatobi. These results suggest that Aceh, Batam-Bitam, and Wakatobi have much greater species retention rates between samples and sites within the region than the other sites.

Species retention rates at the site level showed a greater degree of shared species between all sites than between all replicates samples across all regions except Derawan (Supplemental Figure 19-25). Combined, the above results indicate that for all Eastern regions within the Coral Triangle there was high community turnover between samples and sites with few shared taxa, largely driven by under-sampling at the replicate and site level.

### Comparison eDNA diversity to that of visual census data in Raja Ampat

Comparing eDNA results to visual census survey results from Allen and Erdmann (pers comm.) showed very different patterns across Raja Ampat (Table 2). For all sampling sites with the exception of Kabui Strait, visual census surveys recovered much more fish diversity than total eDNA ASVs. This pattern was more pronounced examining on ASVs identified to species (Table 2). The greatest observed species overlap was in Kri Lagoon where 51 species were detected by both methods for a combined 426 total species. The lowest observed overlap was in Cape Kri with only 17 species shared among a total of 274 identified through eDNA and visual methods. Even though only a limited number of ASVs from eDNA analyses could be identified to species, many ASVs identified to species were not detected in visual surveys.

## Discussion

Environmental DNA (eDNA) is an important metagenomic tool that is effective at assessing community diversity in lower diversity temperate marine ecosystems (Bohmann et al., 2014; Gold et al., 2020; Kelly et al., 2014; Miya et al., 2015; Rees et al., 2017; Thomsen et al., 2012a). Similar to these previous studies, our results highlight the power of eDNA to capture a wide range of fish diversity and detect species that were missed during visual surveys. Importantly, eDNA could distinguish among fish communities across the Indonesian Archipelago, highlighting the distinctiveness of the fish fauna from Aceh that are dominated by Indian Ocean, rather than Pacific Ocean species (Rudi et al., 2012), and Batam-Bintam, a shallow water reef environment on the Sunda Shelf.

However, results also highlight significant challenges for eDNA in regions of exceptional diversity like the Coral Triangle. While eDNA recovered a large amount of fish biodiversity in the form of ASVs, only a fraction of this diversity could be identified to species, highlighting the limitations of existing reference databases, as previously noted for the Coral Triangle biodiversity hotspot (Juhel et al., 2020) as well as lower diversity temperate marine ecosystems (Gold et al., 2020). Another key limitation is that sampling protocols commonly employed in less diverse aquatic ecosystems (e.g., Kelly et al., 2014; Miya et al., 2015; Thomsen et al., 2012a) are insufficient to capture the high fish biodiversity of Indonesian coral reefs. Although rarefaction plots showed that sequencing depth successfully captured all eDNA diversity within individual seawater samples, the total number of samples was inadequate to capture the diversity at any individual site or region. This pattern wasn’t just observed in Raja Ampat, as previously reported by Juhel et al. (2020), but also in lower diversity regions of the Coral Triangle and outside of the Coral Triangle, suggesting the need for refining eDNA sampling approaches in biodiversity hotspots.

### Limitations of Sample Databases

The lack of complete reference sequence databases is a common issue in metabarcoding studies and is noted by Juhel et al. (2020) specifically for Indonesia. They indicate that existing databases cover less than 25% of fish diversity in Raja Ampat, resulting in the identification of only 211 species, with the vast majority of OTUs being unidentified. We improved on this figure by supplementing the *CRUX* database with additional *12S* fish barcodes from the Mo’orea Barcode Project (C. Meyer, pers comm), allowing us to identify a total of 679 species from eDNA, and 55.7% of all ASVs. While an improvement over <25%, this result that highlights the need for expanded barcoding efforts of fish within the Coral Triangle, much like the California Current Large Marine Ecosystem Database developed for California marine fishes (Gold et al., 2020). Developing these resources would allow eDNA to capture and identifying local diversity in biodiversity hotspots as in studies from lower diversity ecosystems (e.g., Andruszkiewicz et al., 2017; Kelly et al., 2014; Miya et al., 2015; Thomsen et al., 2012a), and would provide greater insights into the diversity of fishes being missed by visual surveys.

Surprisingly, one of the lowest percentages of ASV species assignment was in Derawan (57.7%) and Aceh (61.9%) despite supporting much less diverse reef fish communities than Raja Ampat (62.9%) and Lembeh Strait (65.6%) (Allen, 2002). In contrast, only one site had greater than 85% of ASVs identified to species level, Batam-Bintam, the least diverse region sampled. These results highlight two key issues. First, the National Center for Biotechnology Information (NCBI) and the European Molecular Biology Laboratory (EMBL) have a limited number of samples from the Coral Triangle, a function of limited research focus on this region (Barber et al., 2014; Fisher et al., 2011; Keyse et al., 2014); there is no reason to believe that eDNA efforts in the Coral Triangle wouldn’t be as effective if research effort in this region was proportional to diversity. Second, what limited research occurs in the Coral Triangle is often focused on the highest diversity regions like Raja Ampat (Allen & Werner, 2002; Allen, 2008; Roberts et al., 2002), while largely ignoring less diverse regions like Aceh. One reason taxon assignment was as high as it was in Raja Ampat is that a substantial number of Pacific Ocean taxa in habit this region, so the addition of barcodes of fish from Mo’orea increased taxon assignment. However, there are not similar resources for the Indian Ocean or South China Sea. Given the remarkable diversity of the Coral Triangle, it is highly likely that even low diversity regions harbor a substantial amount of unknown biodiversity, suggesting that efforts to develop reference databases should not just focus on regions like Raja Ampat, but should also extend into less diverse and less studied regions.

### eDNA vs Visual Census

A common feature of marine eDNA studies is their ability to return more diversity than traditional survey methods (Kelly et al., 2017; Stat et al., 2017, 2019). However, in all but one case in this study, eDNA recovered substantially less fish biodiversity than visual census data from the same location. In side-by-side comparisons in Raja Ampat, not only did eDNA recover much less fish diversity than visual census methods, but it also recovered completely different fish diversity. At most, only ~17.4% of taxa overlapped in eDNA and visual census surveys (Table 2).

While this result, in some ways, highlights the challenge of eDNA approaches in megadiverse ecosystems, it also highlights its utility. In total, eDNA recovered an additional 168 species of marine fishes spanning 59 families and 125 genera that were missed by visual census surveys. While some of the diversity captured by eDNA included cryptic taxa like Blennidae that are easily missed in visual census surveys, it also captured a large number of taxa like Lutjanids, Pomacentrids, Siganids, and Epinephalids, among others, that are large, conspicuous fish that should be easily observed.

While it is possible that eDNA at any given location could have been transported from a nearby reef, or represent eDNA from larval stages or eggs, the comparison of taxa identified through eDNA and visual census was done at the regional, not local scale minimizing this concern. Moreover, phylogeographic analyses show that Raja Ampat and Eastern Indonesia are phylogenetically and biogeographically very unique (Barber et al., 2011; DeBoer et al., 2014) due to geology and physical oceanographic processes (Kool et al., 2011). As such, it is unlikely that eDNA would be transported from a different biogeographic region. However, because of the phylogeographic uniqueness of Eastern Indonesia (Carpenter et al 2010, Barber et al. 2011), it is possible that sequences representing a particular species obtained from eDNA are genetically divergent from sequences in public reference databases, and that a portion of the unique ASVs reflect deep phylogeographic structure, such as that observed in other marine taxa (Adams et al., 2019). Combined, these results highlight the complementary nature of eDNA to more traditional fish biodiversity survey methods (Stat et al., 2019).

### Importance of Sampling Intensity

The results of this study are likely a function of sampling intensity. Visual census of fish communities on coral reefs are done in multiple habitats over multiple days to maximize the amount of diversity recovered (Allen, 2002; Erdmann & Pet, 2002). While we sampled multiple liters of sea water from across individual reefs and from multiple reefs within a given region, visual census data required multiple divers spending 6hrs/day for 10-21 days for ****each**** of the regions we sampled in Raja Ampat. Given that visual sampling dwarfed our sampling design, it is perhaps unsurprising that eDNA recovered much less diversity.

However, results also suggest limits to the diversity eDNA can recover. One striking pattern revealed in this study is that despite greatly under sampling total fish diversity, we achieved saturation in sequencing depth for every individual sample. Thus, despite the Coral Triangle having levels of teleost diversity that dwarfs Southern California kelp forests, the latter has average levels of per sample fish diversity similar to the Coral Triangle (Figure 2). This unexpected result suggests that there may be an upper limit to the amount of diversity eDNA can capture within a single liter of seawater. As such, eDNA studies in high diversity ecosystems will require much greater sampling intensity, a result consistent with Juhel et al. (2020). Given that our models indicate that high biodiversity regions like Raja Ampat may require in excess of 300 one-liter samples to capture all fish diversity present, eDNA surveys in high diversity areas will either need to increase the numbers of samples or volume of water per sample to be maximally informative. Further studies are necessary to determine the optimal sampling strategy and potentially novel developments in eDNA metabarcoding including massively multiplexed pooling methods combined with even higher throughput sequencers (Singer et al., 2019).

We also highlight the importance of using a zeta diversity framework for exploring patterns of biodiversity through eDNA metabarcoding analyses (Lin et al. 2020). Here zeta diversity allowed us to disentangle the nested nature of eDNA sampling schemes and explore community turnover across samples, sites, and regions. Here we found stark rates of zeta decay across most sites sampled (Figure 5). Intuitively, we expect the shared diversity to decrease as you increase the number of communities observed since fewer species are likely to be found across a greater number of sites (Hui & McGeoch, 2014). This relationship has been well characterized both theoretically and empirically (Hui et al., 2014; McGeoch et al., 2019). Importantly, in areas with higher biodiversity and rates of community turnover we expect fewer species to be shared between communities and thus we expect to observe a faster rate of zeta decay. Here we showed that evaluating the rates of zeta decay across samples and sites separately allowed us to disentangle the higher community overlap in less diverse Western Indonesia sites (Ache and Batam-Bintang) from the severe under sampling of coral reef fish diversity given our sampling design.

Similarly, species retention rates thus provide important information on the rarity of species shared across communities (Latombe et al., 2018). In areas with a few common taxa across all communities, species retention rates in less diverse communities will saturate faster than in more diverse communities as the few common taxa comprise a greater proportion of species within the community (Krasnov et al., 2020). In contrast, in high biodiverse regions with very high rates of community turnover, we expect a more bell-shaped curve in which species retention rates initially increase as rare taxa are lost across a few sampled communities but then rapidly decreases as very few species or none at all are present in more than a handful of observed communities (Latombe et al., 2018). Here we found that lower diverse Western sites saturated much quicker than the higher diverse Eastern sites (Raja Ampat and Lembeh Strait) at the site level, indicating that community turnover between sites was much greater in the center of the Coral Triangle as expected. Across replicate one liter samples, we similarly found that sites within the heart of the Coral triangle had lower species retention rates and small proportion of shared common taxa across all sites. Interestingly, the bell shaped species retention curve observed in Batam Bintam suggests low community overlap between samples despite having the highest species retention rates across sites. However, combined with the low zeta decay results above this is a function of many shared species within the three replicate samples at each site in contrast to very few shared taxa across sample replicates observed at all other sites. Thus, combining species retention rates and zeta decay allows for the comparison of community turnover rates between biogeographic regions and provides a more complete characterization of the patterns of biodiversity between communities observed through multiple eDNA samples and sites.

### Biodiversity patterns across Indonesia

One of the best-documented patterns of marine biodiversity is the extreme concentration of biodiversity in the Coral Triangle (Bellwood & Meyer, 2009; Roberts, 2002). Given that our study stretched from regions like Raja Ampat, known for the highest marine fish diversity in the world (Allen, 2002, 2008) to regions in Western Indonesia that are outside of the Coral Triangle (Veron et al., 2009), and given that these patterns are based partially on fish biodiversity (Bellwood & Meyer, 2009; Roberts et al., 2002), we expected to see clear gradients in diversity based on eDNA. However, while the Eastern Indonesian reefs of Raja Ampat had the highest fish biodiversity, and the Western Indonesian reefs of Batam-Bintam had the lowest, there was no clear gradient in between.

There are four potential explanations for the failure of eDNA to capture this east-west biodiversity gradient in Indonesia. First, the rarefaction plots and models of sampling depth required to achieve ASV saturation show that eDNA recovered a higher percentage of fish diversity present in Western Indonesia than in Eastern Indonesia. Thus, the inability to recover a well-known biodiversity pattern could be an artifact of severely under-sampling the most biodiverse reefs in Indonesia, and that higher intensity eDNA sampling design could yield results that conform to predictions based on previous studies.

A second, but not mutually exclusive explanation, is that visual surveys may not fully capture all of the diversity present. There are many taxa like blennies and gobies that live within the reef matrix (Böhm & Hoeksema, 2017; Kotrschal, 1988; Nursall, 1977; Wilson, Fisher, & Pratchett, 2013). Moreover, visual surveys in Indonesia have been done on open-circuit SCUBA, which can negatively impact fish counts (Lindfield et al., 2014). As such, the well-known biodiversity gradients in Indonesia may reflect visually conspicuous biodiversity, but not cryptic diversity or fish diversity sensitive to SCUBA.

A third potential explanation to the mismatch between eDNA and visual census data is that these studies were not conducted contemporaneously. Given that marine ecosystems are dynamic, conducting eDNA and visual surveys at different times could result in capturing different communities (Beenties et al., 2019). This possibility would be easily tested by including eDNA sampling as part of any fish visual census protocol (Alexander et al., 2020) and doing so could add value to visual surveys as eDNA may provide insight into parts of the fish community (e.g., blennies) that are difficult to fully capture in visual surveys (Alexander et al., 2020; Stat et al., 2019).

Finally, because eDNA and visual census survey data showed minimal taxonomic overlap it is possible that the more cryptic diversity captured by eDNA does not follow expected regional patterns in biodiversity, a result that could be confirmed through studies of cryptic invertebrate taxa or fish surveys on rebreathers, rather than open-circuit SCUBA. Given the above, combined with the fact that 1) Aceh is located at the nexus between the Indian Ocean and Java Sea (Fadli et al., 2012, 2014), 2) this region is a suture zone between Pacific and Indian Ocean basins (Crandall et al., 2012), and 3) this region is receiving less scientific study thatn regions like Raja Ampat, it is possible that Western Indonesia isn’t as biologically depauperate as presently believed.

### Prospects for eDNA in high diversity ecosystems

While eDNA did not perfectly capture patterns of Indonesian fish biodiversity, results broadly conform to expectations. While eDNA did not perform as expected in comparison to visual surveys, it is important to put these numbers into the context of sampling effort and cost—eDNA sampling was done quickly and inexpensively without decades of taxonomic expertise, while visual surveys were expensive, as well as time and labor intensive. As such, eDNA clearly has value in high diversity ecosystems, despite its limitations. In particular eDNA can 1) provide substantial insights into marine biodiversity with minimal taxonomic knowledge and, 2) provide complementary data on rare or cryptic taxa potentially overlooked by traditional visual surveys, highlighting the immense value and promise of eDNA data.

eDNA is clearly a valuable new tool for examining biodiversity across a variety of habitats (Closek et al., 2019; Jeunen et al., 2019; Port et al., 2016; Stat et al., 2017; Stoeckle, et al., 2020). As the field of eDNA continues to mature, it is critical to better understand the performance of eDNA in highly diverse ecosystems, such as Indonesia (Juhel et al., 2020). Important next steps include working to develope reference databases in diverse tropical marine ecosystems like the Coral Triangle to increase the number of ASVs that can be assigned to species (Ward et al., 2009). In addition, performing eDNA and visual surveys at the same time in the same location, and increasing either the volume or number of eDNA water samples, could provide important insights into the best sampling design for eDNA research, creating standard field protocols for eDNA and visual census surveys in the future (Deiner et al., 2017).

## Acknowledgements

We thank the Indonesian government, including the Ministry for Research Education and Technology and the Indonesian Institute of Sciences for research permits to conduct this study (SK Kepala Lembaga Ilmu Pengetahuan Indonesia No. 1834/H/2018). This work was funded by USAID PRESTASI Scholarship Program and the National Science Foundation (OISE 1243541) and was a collaboration research between the Research Center for Oceanography, Indonesia Institute of Sciences (RCO – LIPI) and UCLA. Thanks to the late I.S. Arlyza for assistance with sample collection, G.R Allen and M.V. Erdmann who shared data on fish diversity data, and the Coral Reef Information and Training Center – Coral Reef Rehabilitation and Management Program – Coral Triangle Initiative – Indonesian Institute for Sciences (CRITIC COREMAP CTI LIPI) for data sharing on marine fish diversity. M. Hafizt (RCO-LIPI) assisted with map generation.

---

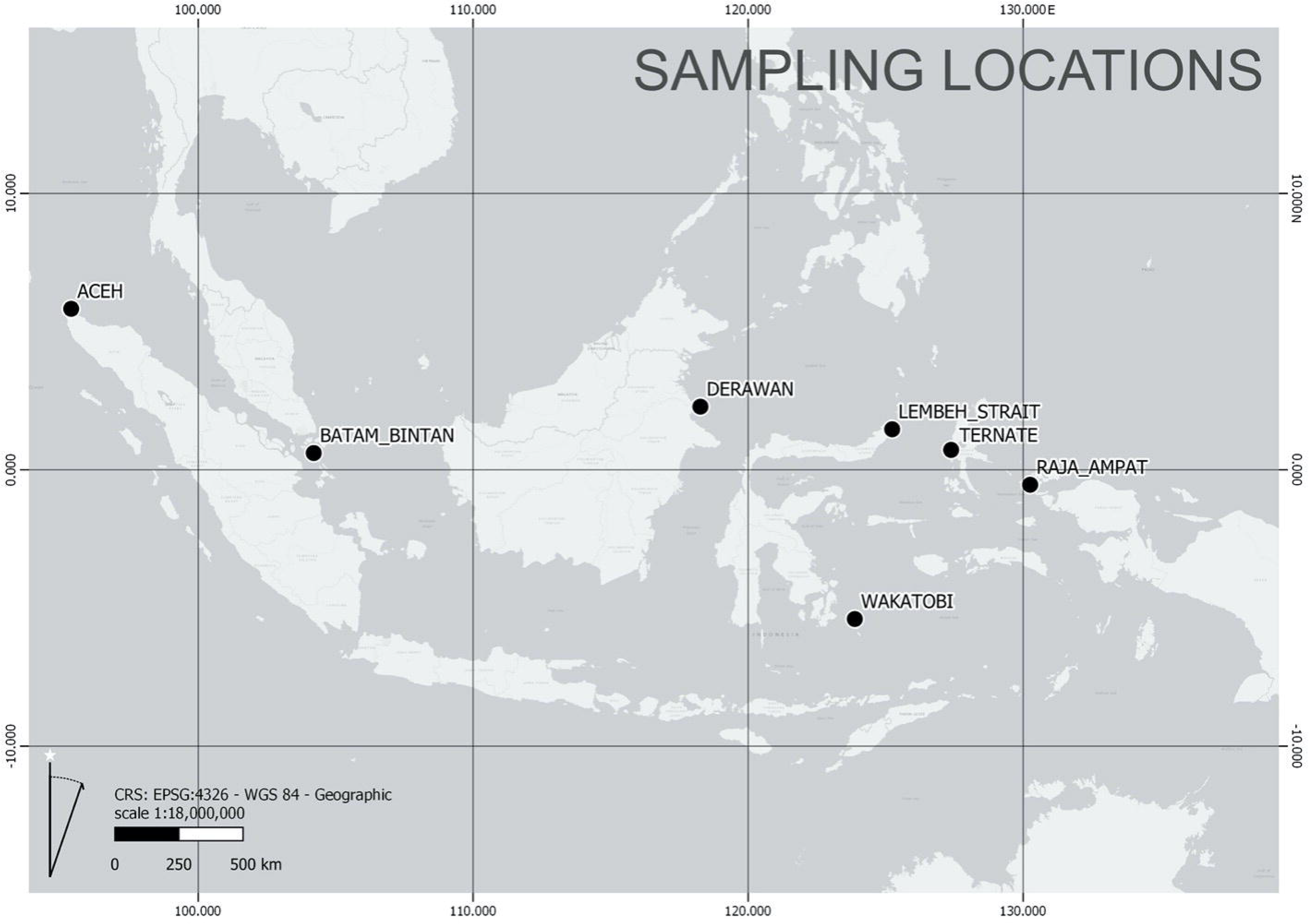

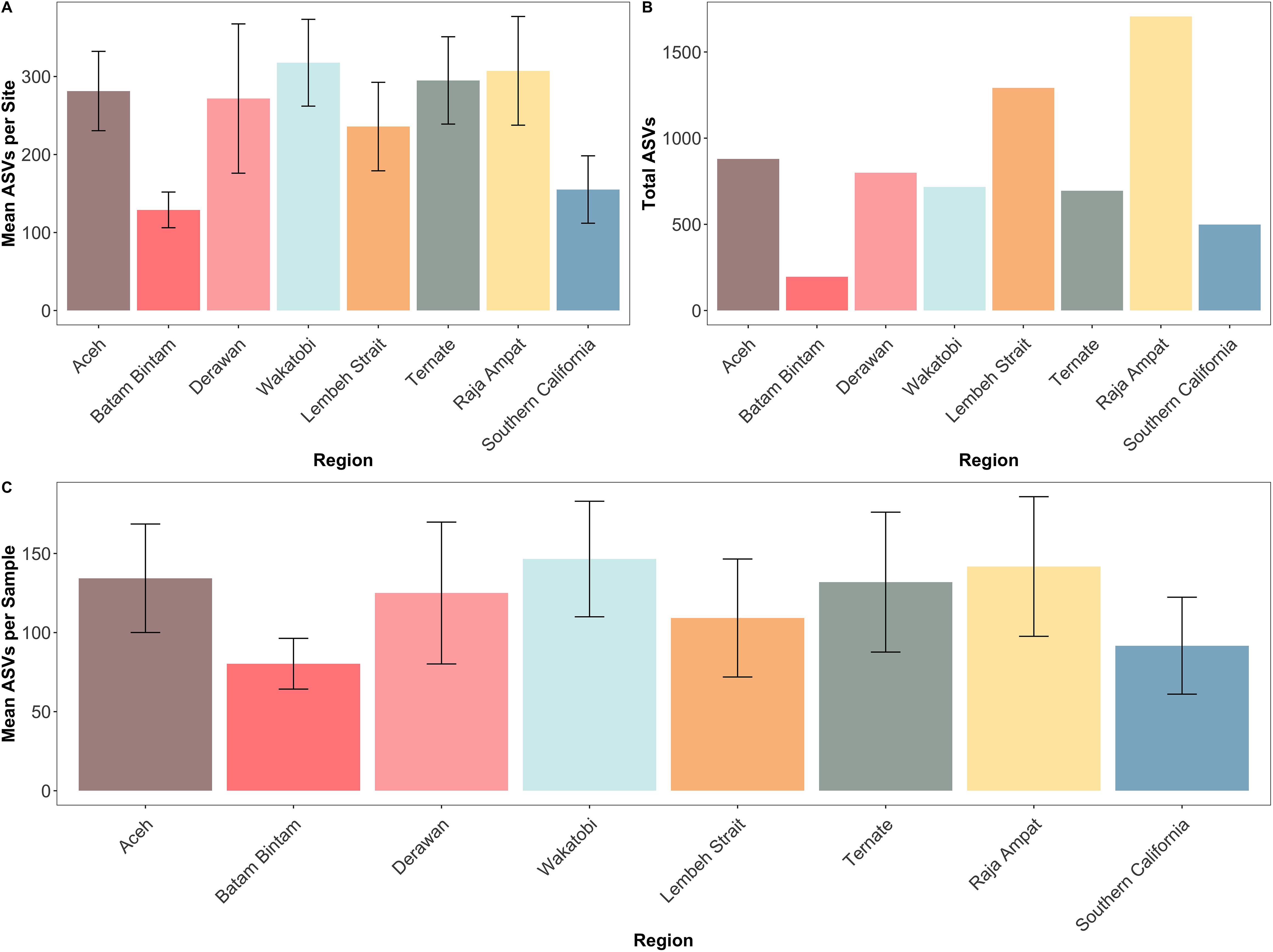

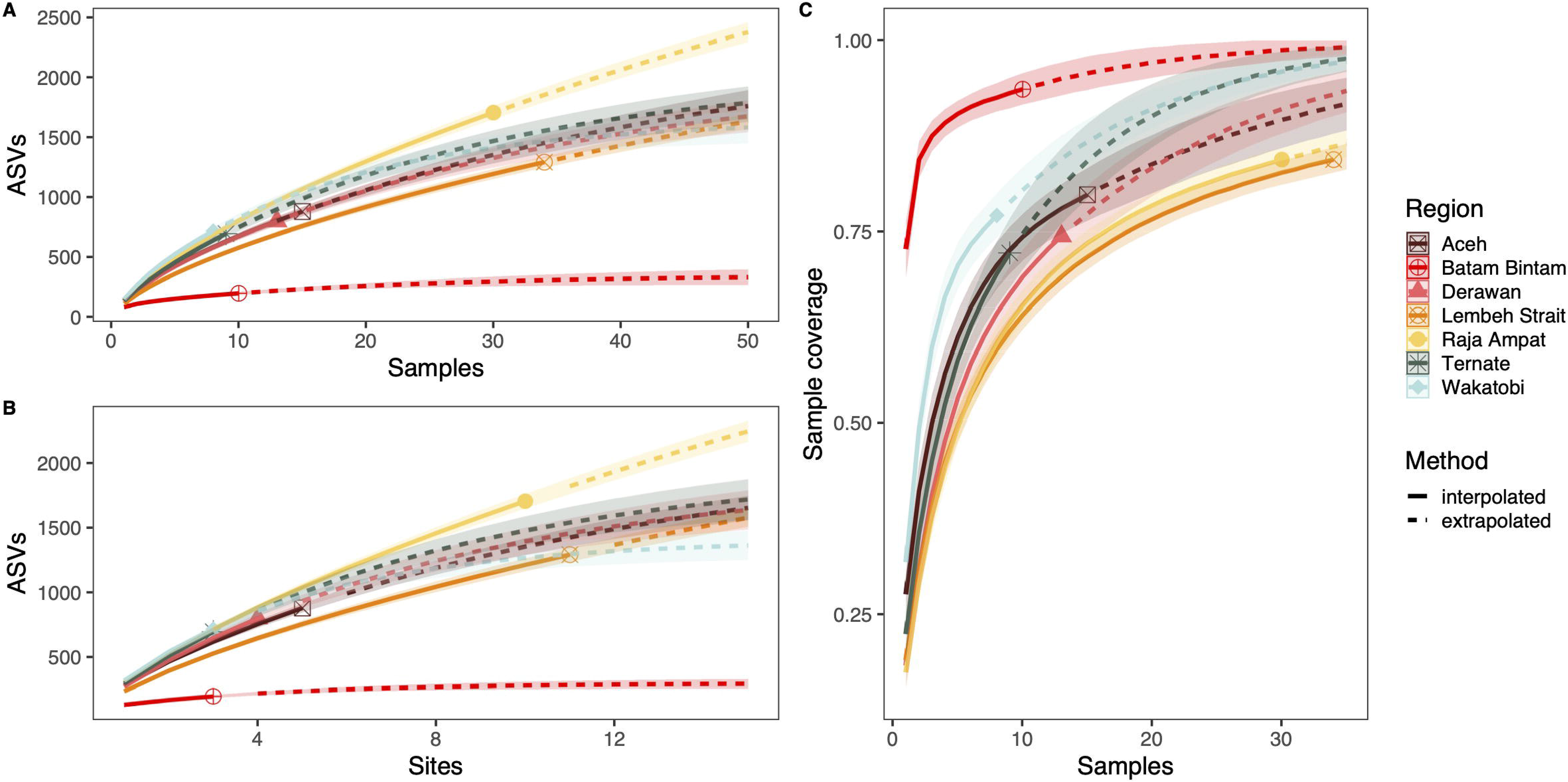

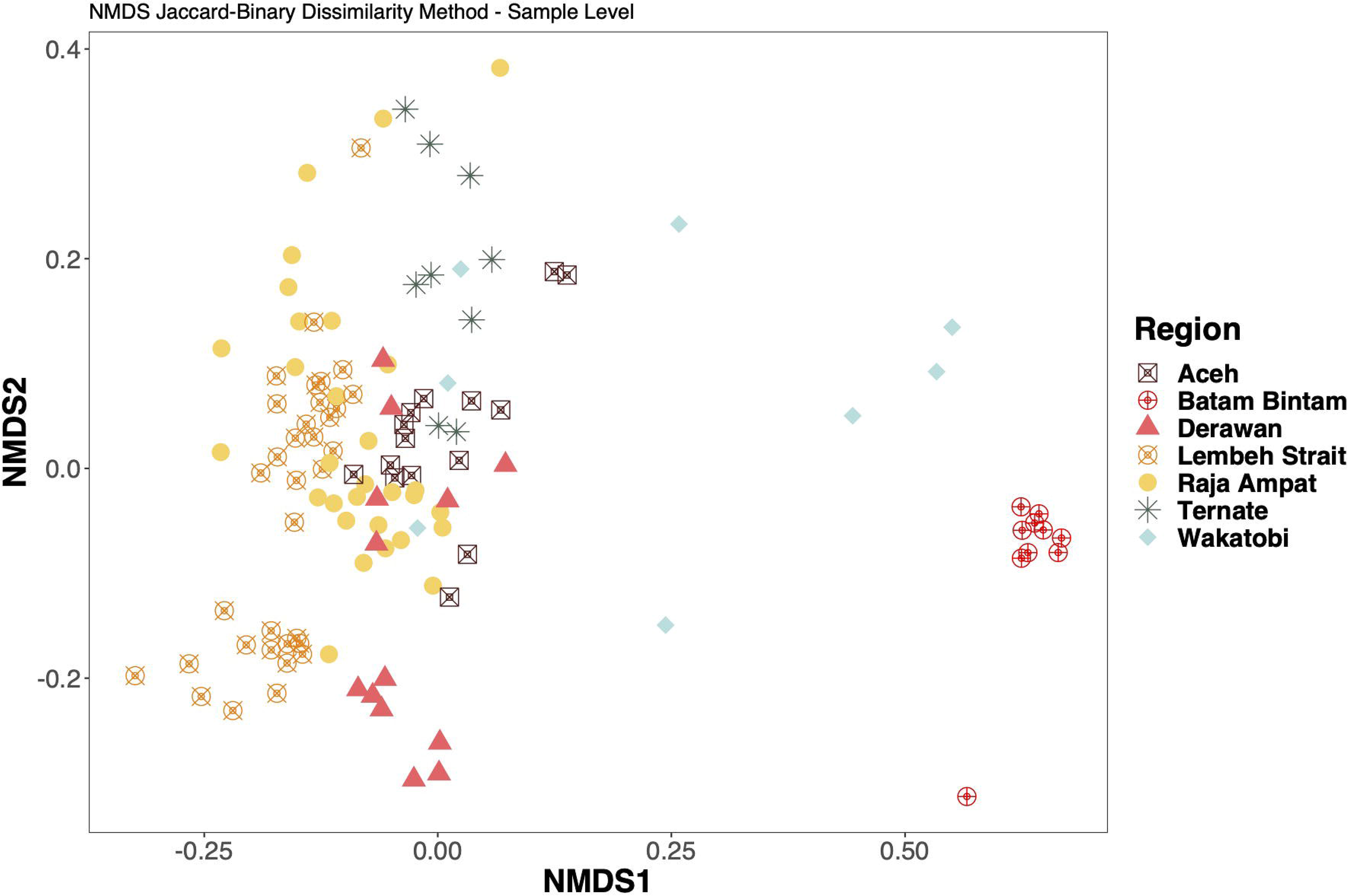

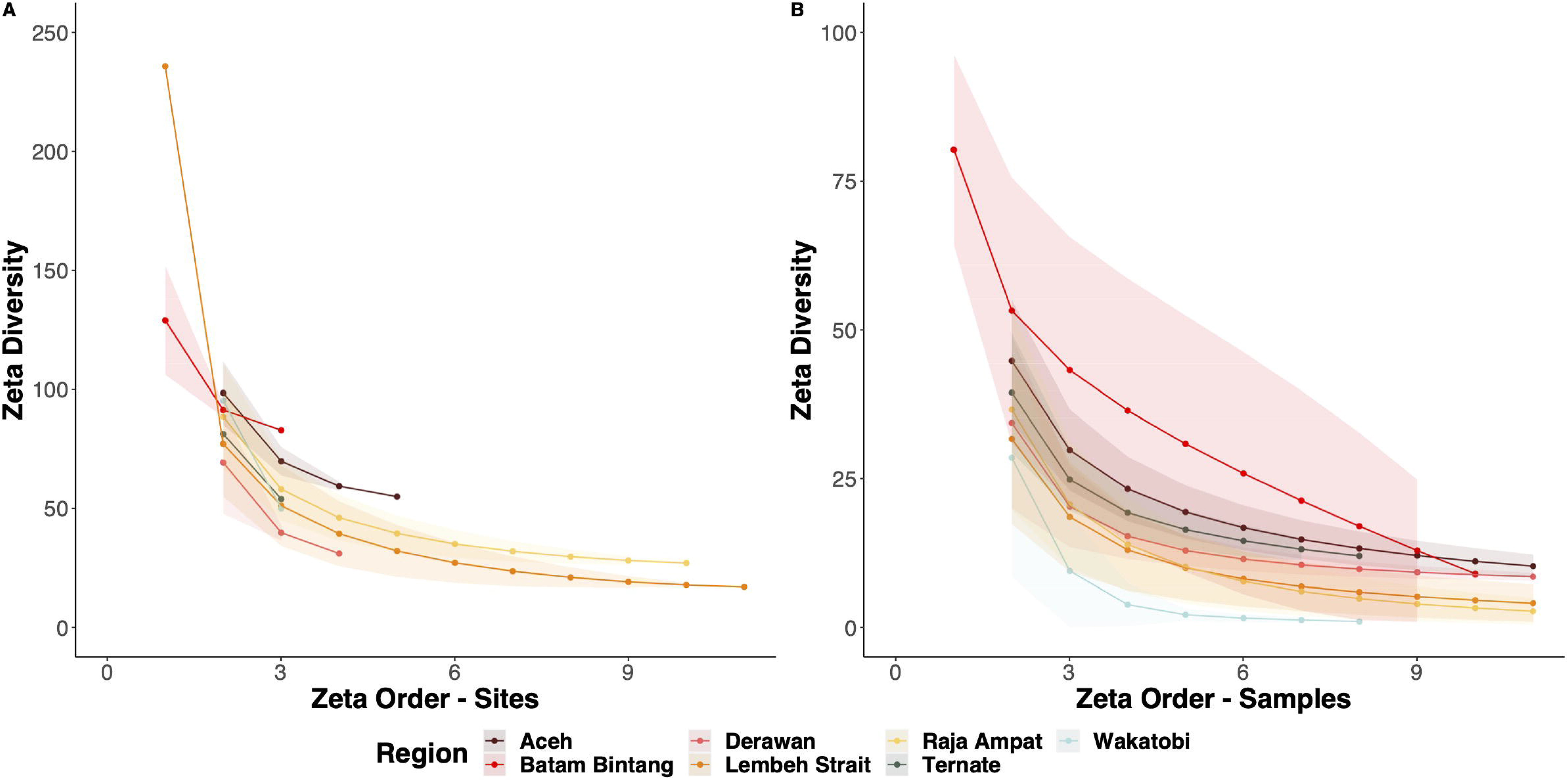

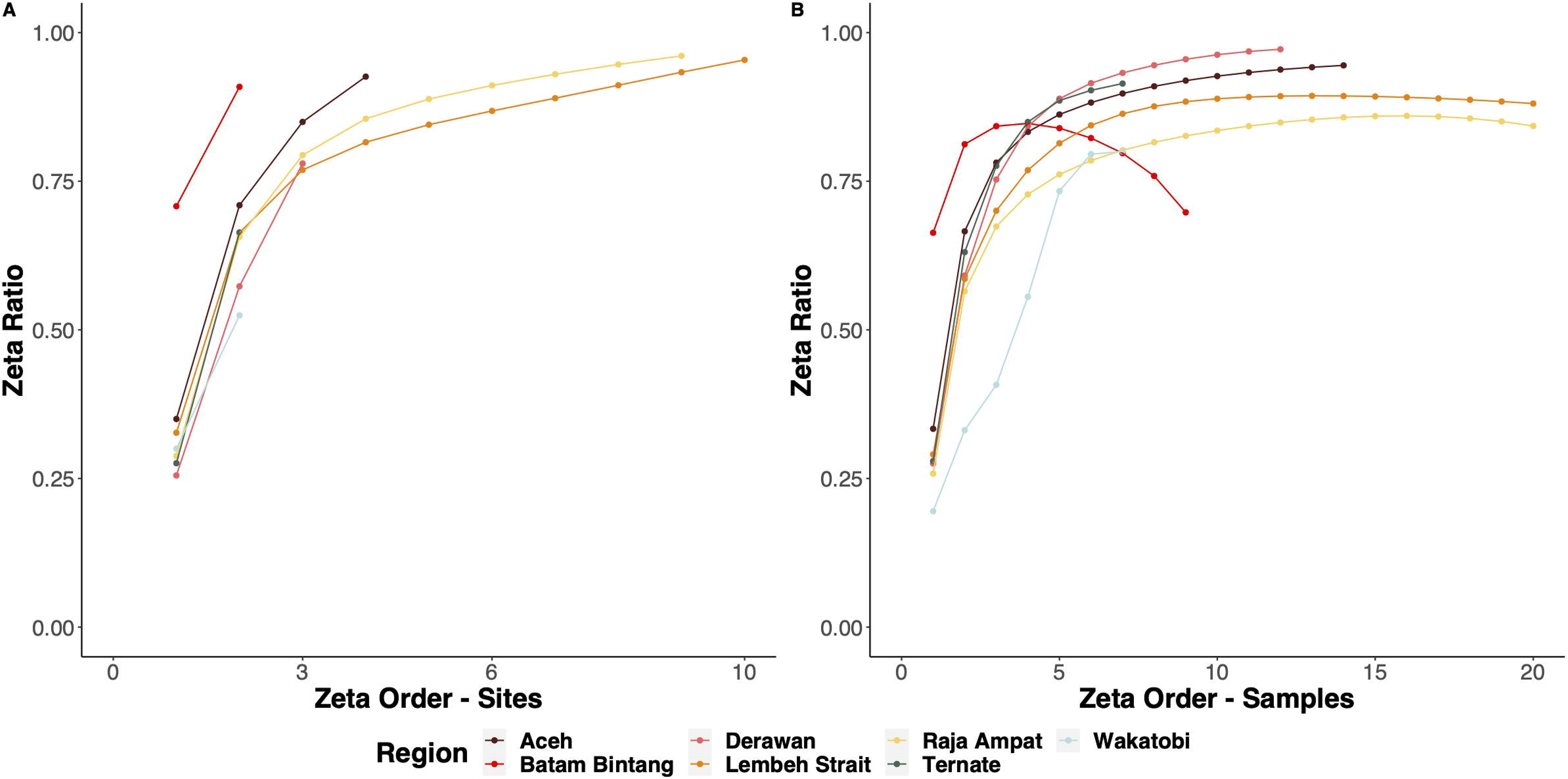

## Notes

### Competing Interest Statement

The authors have declared no competing interest.

