## Supplemental Figures for "Environmental DNA in a Global Biodiversity Hotspot: Lessons from Coral Reef Fish Diversity Across the Indonesian Archipelago"

**Supplemental Information**

**Supplemental Methods**

**Decontamination**

The following decontamination pipeline is an adaptation of the Gruinard decon decontamination pipeline as described in Gold et al. 2020 . This framework builds off of the decontamination scripts outlines in Kelly et al. 2018 and McKnight et al. 2019. Importantly, we did not conduct site occupancy modeling as our goal was to conduct ASV accumulation curves which are sensitive to the tail end distribution of rare ASVs.

*Removing Single End Read ASVs and Samples*

We filtered out all non-paired end reads (forward, reverse, and unmerged paired reads) retaining only paired merged reads. We also removed all samples with less than 5,000 total reads per barcode.

*Estimation of Index Hopping*

All samples were pooled into a final library and sequenced on two MiSeq runs, one per barcode. Each sample is identified by two sets of molecular barcodes in a unique combination. However, recent evidence has found that there is the potential for indexes to hop from one molecular to another, leading to the incorrect sample assignment during demultiplexing (Costello et al., 2018). To estimate the frequency of index hopping we included a positive control of a non-native fish taxa which we know will not be found in our eDNA samples (Kelly, Gallego, & Jacobs-Palmer, 2018). Index hopping will lead to environmental sequences occurring in the positive control and vice versa. To estimate the frequency of index hopping we modeled the composition of environmental sequences observed on the positive controls and subtract these sequences from the environmental samples run. For example, if 12 reads of Garibaldi (*Hypsypops rubicundus*) are found in the positive control, we subtract 12 reads from the read counts of Garibaldi found in all environmental samples.

*Remove Contamination from Negative Controls*

Here we remove ASVs that occur in positive and negative controls with higher proportions than environmental samples using *R* package *microDecon* (McKnight et al., 2019). We used the standard parameters and grouped samples by location.

*Calculating eDNA Index Scores*

The eDNA index was computed following the methods of Kelly et al. 2019 (Kelly, Shelton, & Gallego, 2019). This was accomplished by first calculating the mean read count for each assigned taxonomy and then calculating the relative abundance of each ASV; number of reads of each ASV divided by the total number of reads per sample. The relative abundance of each taxa in each sample was then divided by the maximum abundance for a given species across all samples to generate the eDNA index. The index thus normalizes the read count per species and per sample. The eDNA index values 0 to 1 for each taxa, allowing for abundance comparisons of a specific taxa across sites.

*Removing Highly Divergent Replicates*

Sometimes a sample replicate will amplify and sequence successfully, but the result is clearly due to contamination or off target sequencing as the sample is dominated by a handful of species not present in other samples, or negative and positive controls. An example would be a black fly lands on the lid of at tube and the resultant sample comes back as nearly 100% black fly. This can also occur in samples with low sequencing depth and poor ASV saturation. This typically manifests as a sample replicate having high dissimilarity to other replicates. In this case we used a dissimilarity threshold of >99.9% to remove any one liter bottle replicates that were extremely divergent from each other. Dissimilarities between sample replicates were calculated using the Bray-Curtis dissimilarity index on eDNA index transformed reads.

**Supplemental Figures**

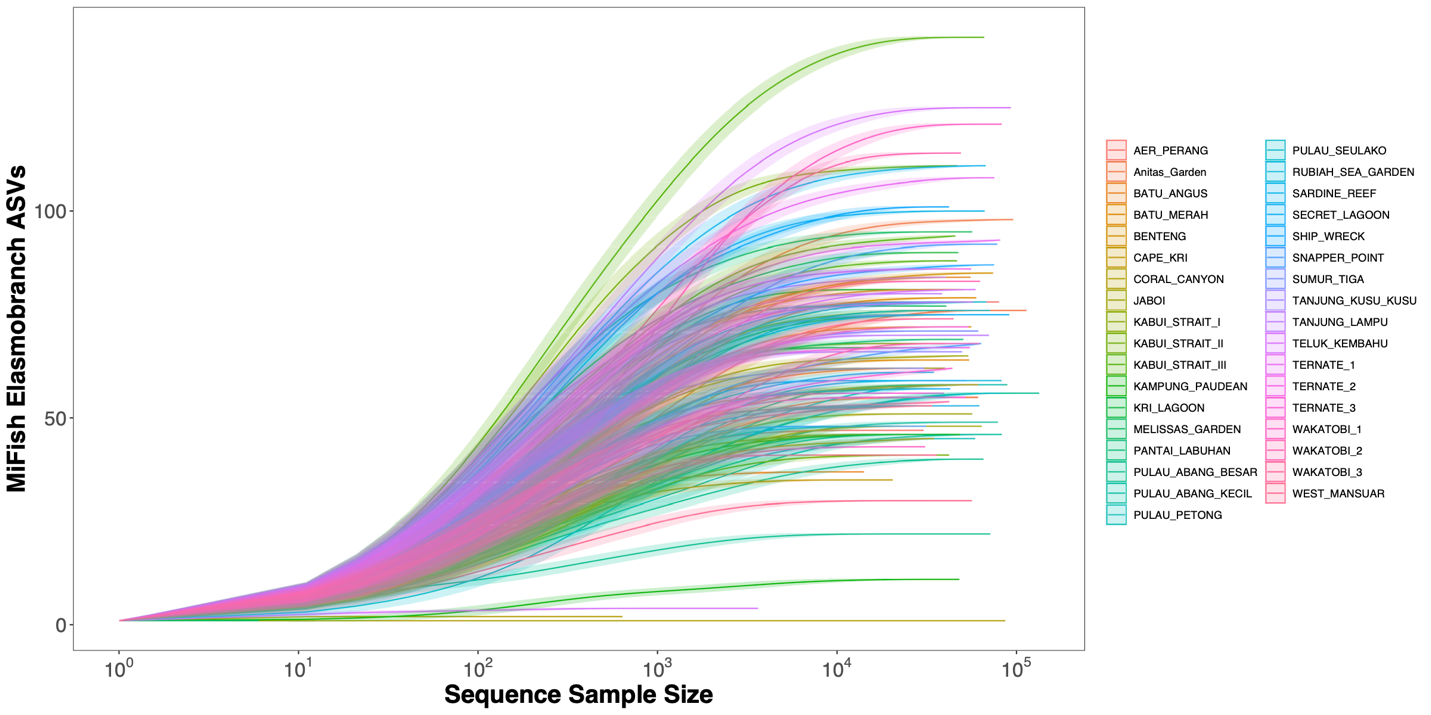

**Figure S1. MiFish Elasmobranch ASV Sequence Rarefaction Curve.**

Individual one liter samples are colored by site locality.

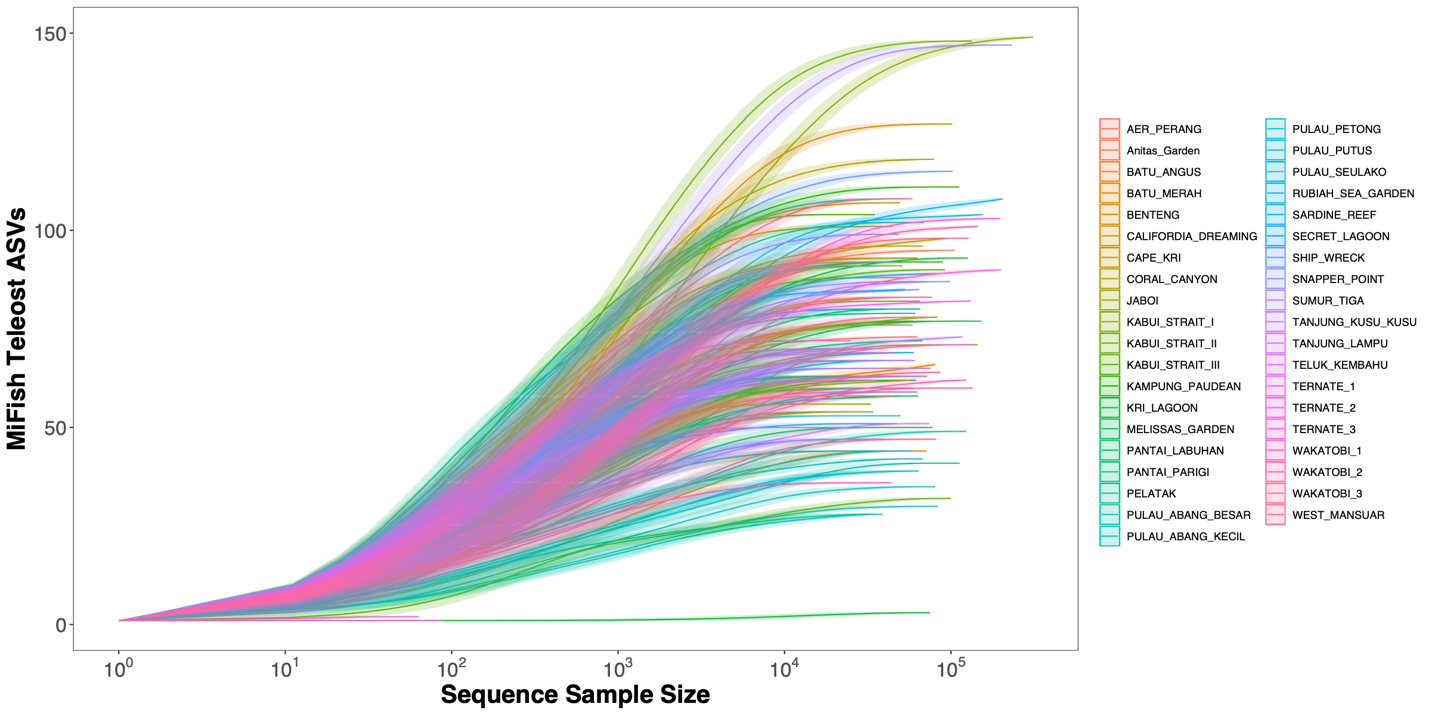

**Figure S2. MiFish Teleost ASV Sequence Rarefaction Curve.**

Individual one liter samples are colored by site locality.

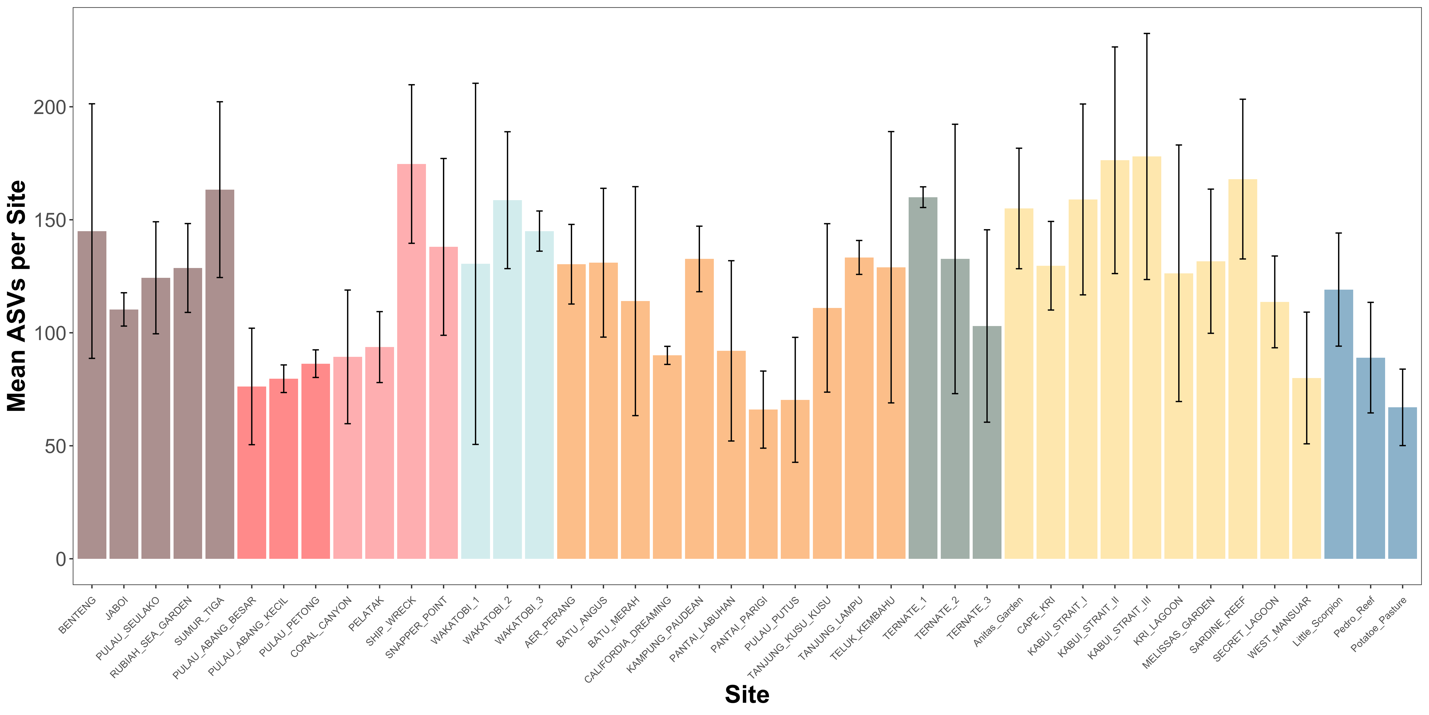

**Figure S3.** Mean Amplicon Sequence Variant (ASVs) richness per Site

**Figure S4. Aceh ASV Accumulation Curve.**

Colored by site locality.

**Figure S5. Wakatobi ASV Accumulation Curve.**

Colored by site locality.

**Figure S6. Batam Bintang ASV Accumulation Curve.**

Colored by site locality.

**Figure S7. Ternate ASV Accumulation Curve.**

Colored by site locality.

**Figure S8. Derawan ASV Accumulation Curve.**

Colored by site locality.

**Figure S9. Lembeh Strait ASV Accumulation Curve.**

Colored by site locality.

**Figure S10. Raja Ampat ASV Accumulation Curve.**

Colored by site locality.

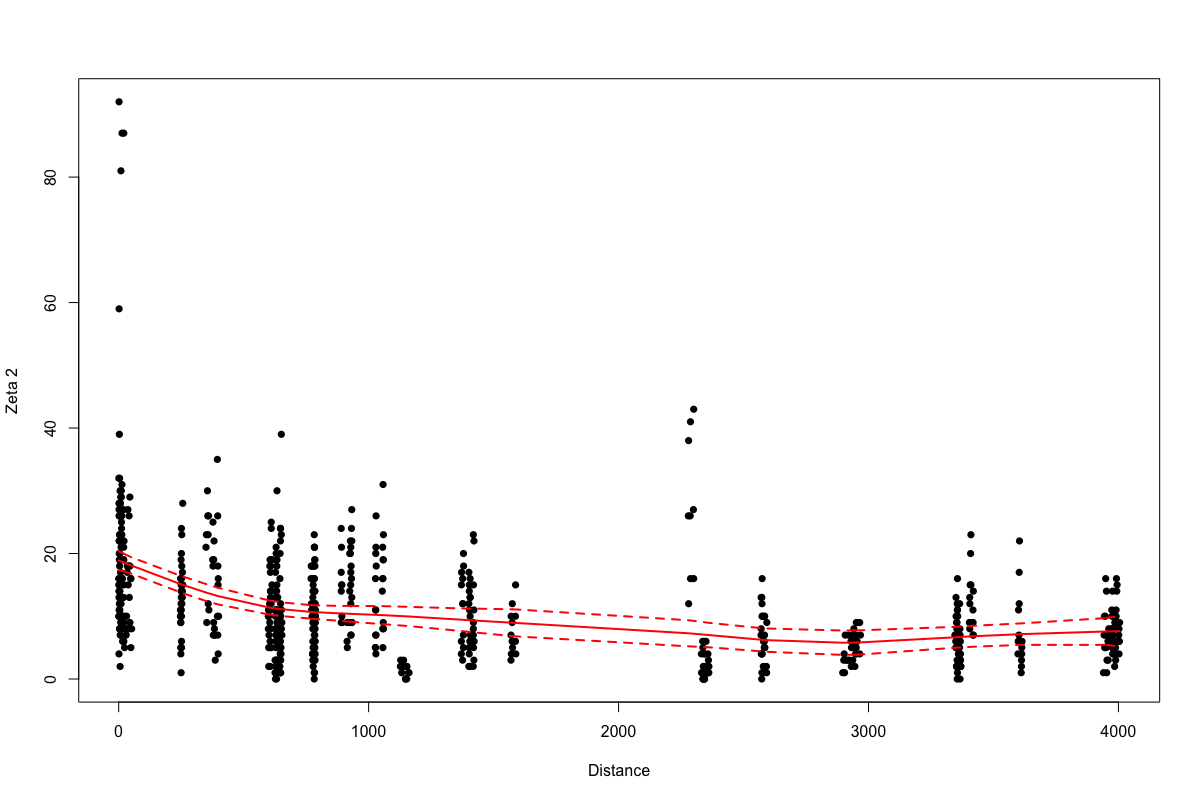

**Figure S11. Zeta Decay Over Distance.**

Plot of pairwise (zeta order 2) ASV diversity across distance. Samples taken from closer proximity (less than 10 km) had greater shared ASVs than those taken from further away.

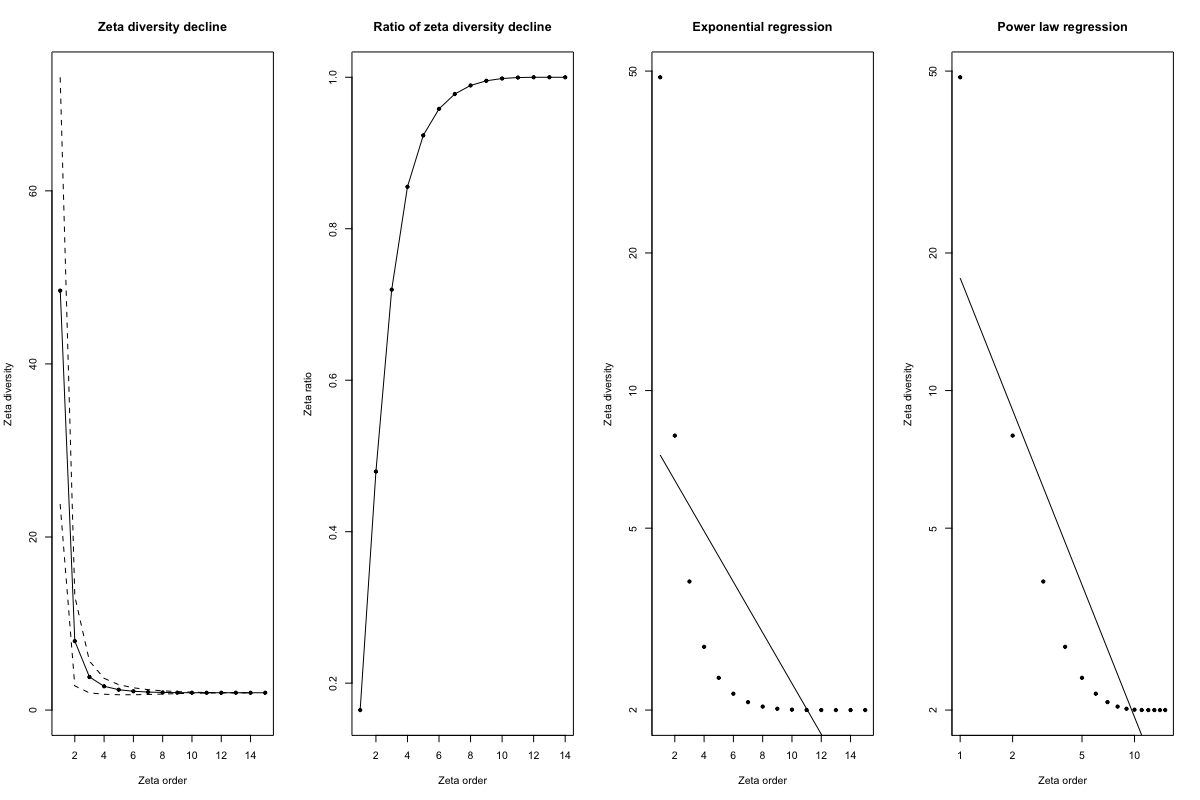

**Figure S12. Zeta Decay of Samples from Aceh.**

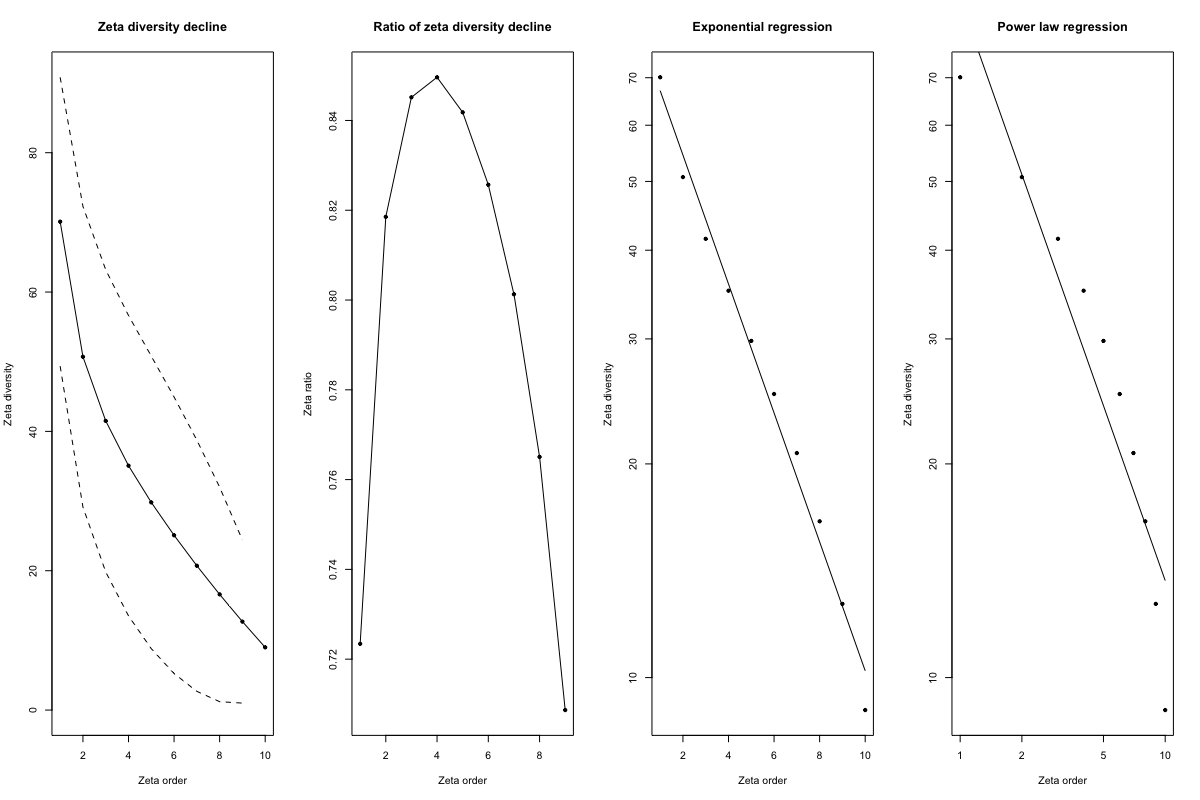

**Figure S13. Zeta Decay of Samples from Bantam-Bintang.**

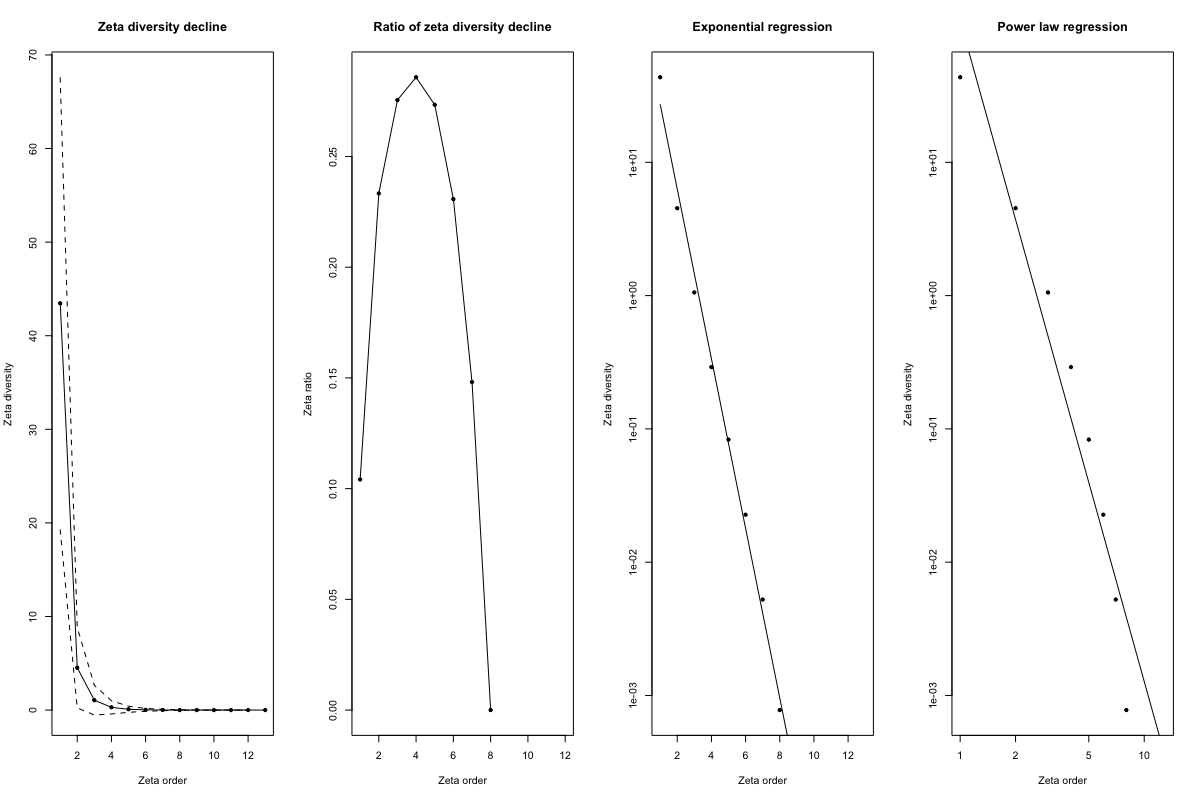

**Figure S14. Zeta Decay of Samples from Derawan.**

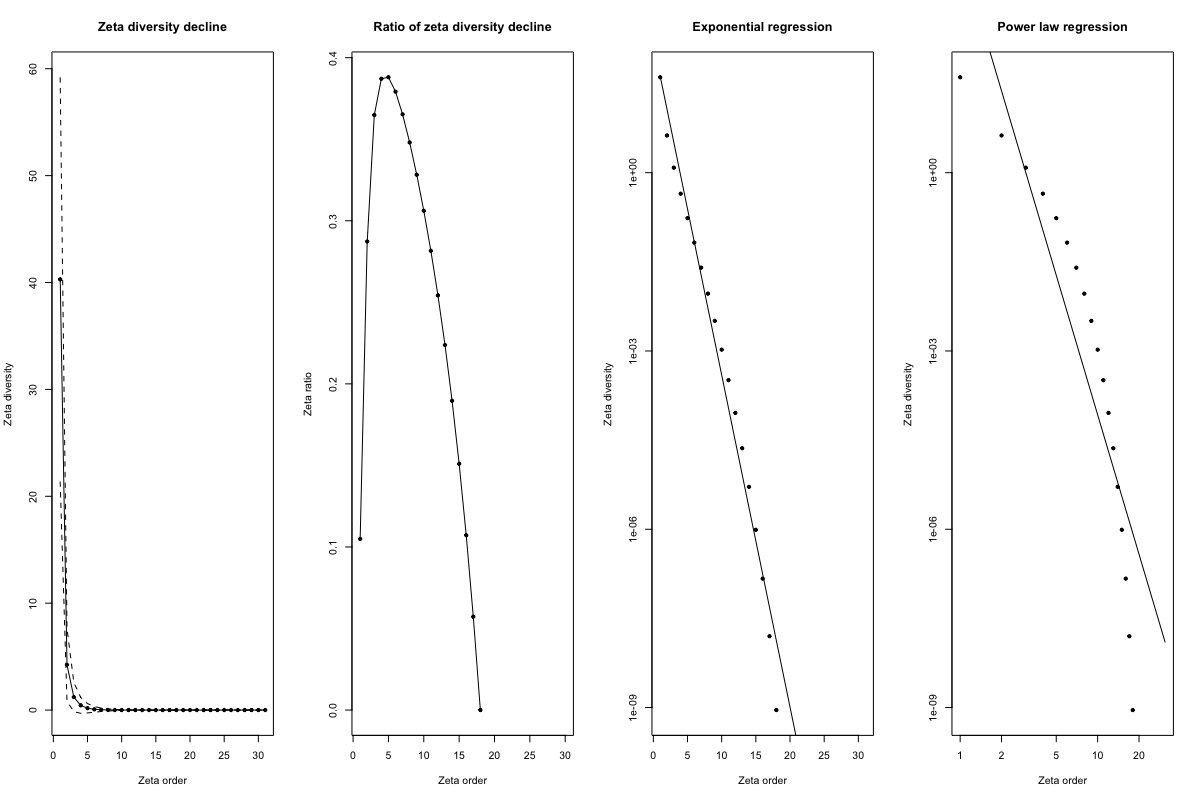

**Figure S15. Zeta Decay of Samples from Lembeh Strait.**

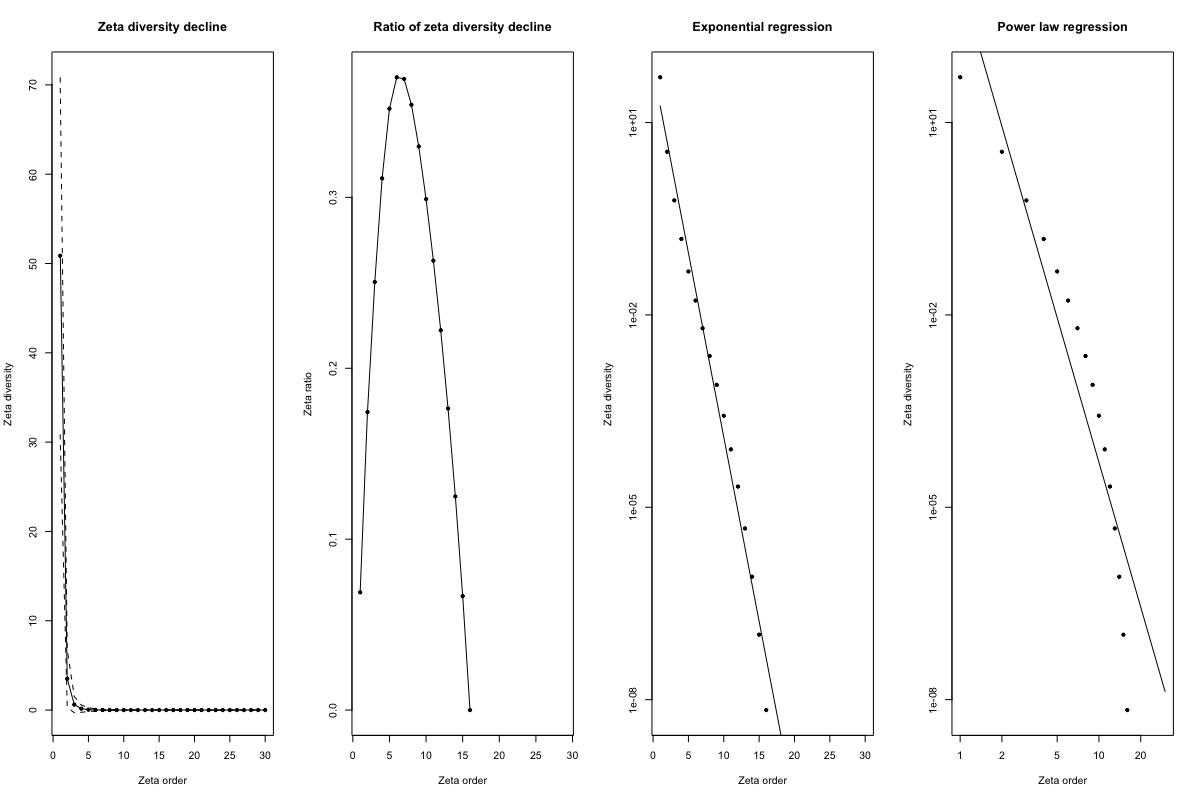

**Figure S16. Zeta Decay of Samples from Raja Ampat.**

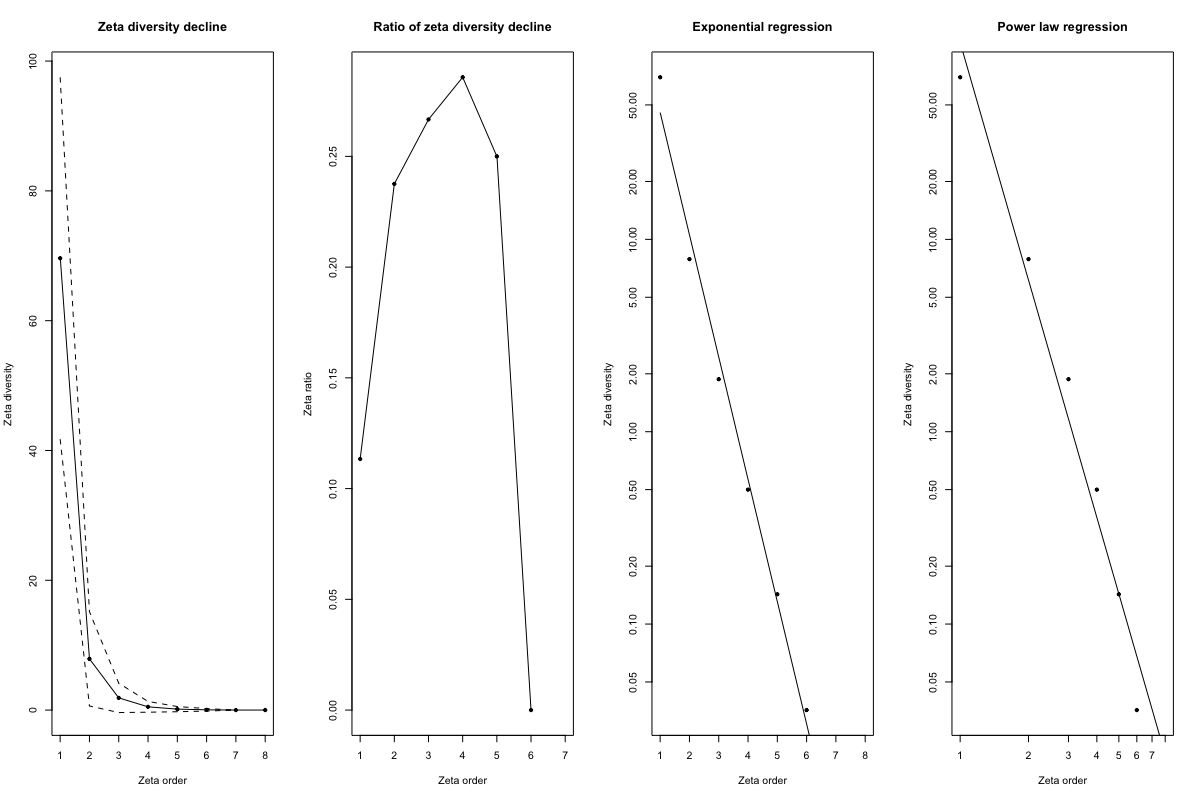

**Figure S17. Zeta Decay of Samples from Ternate.**

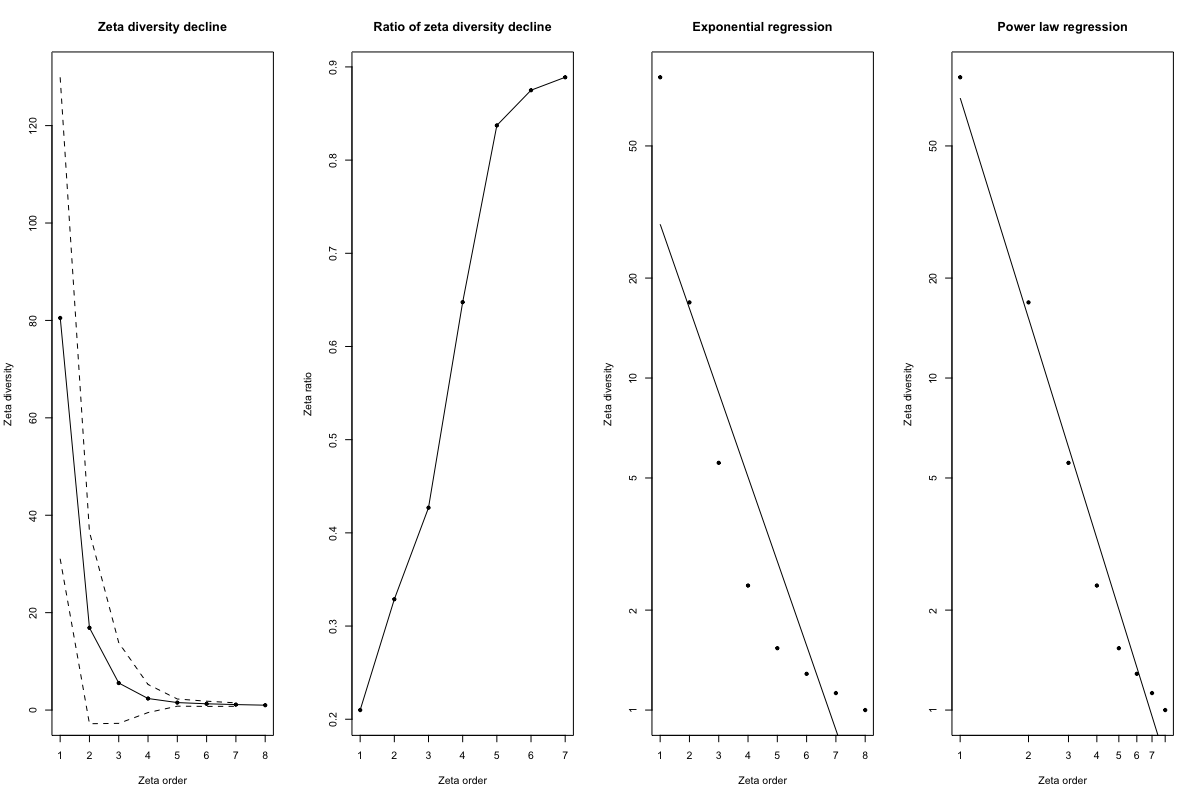

**Figure S18. Zeta Decay of Samples from Wakatobi.**

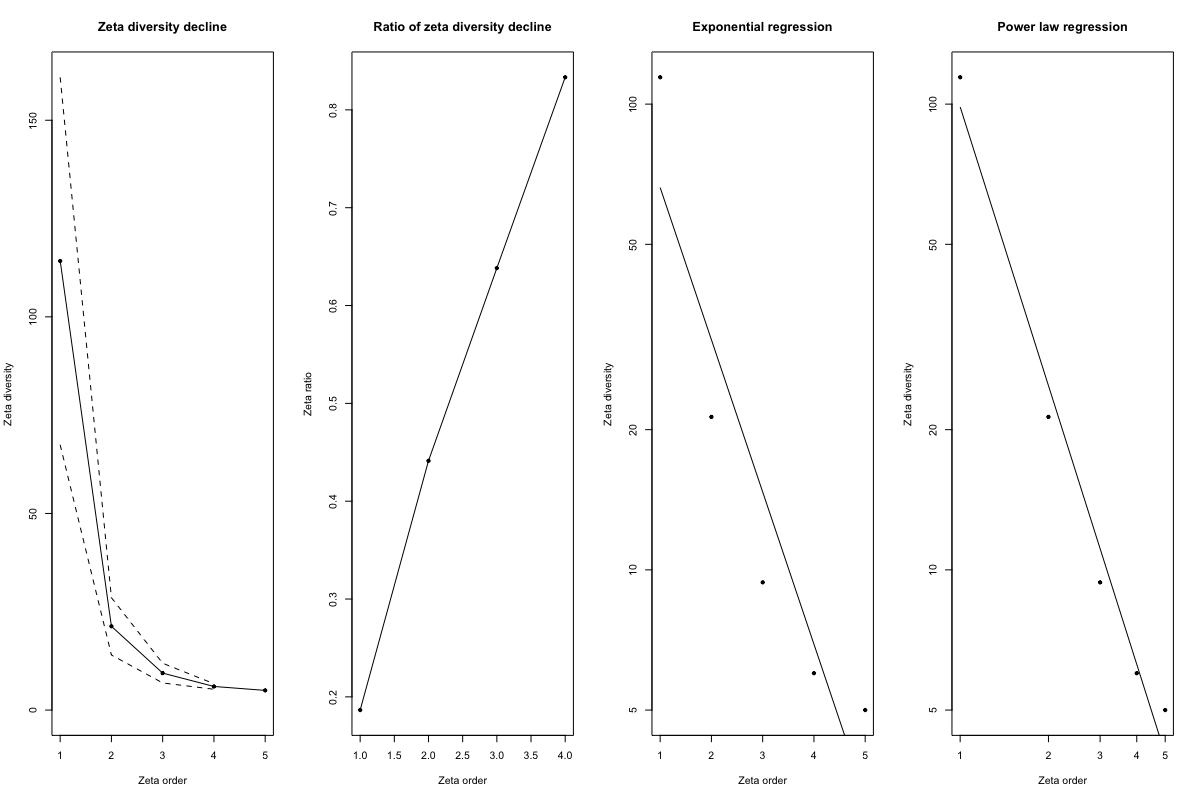

**Figure S19. Zeta Decay of Sites from Aceh.**

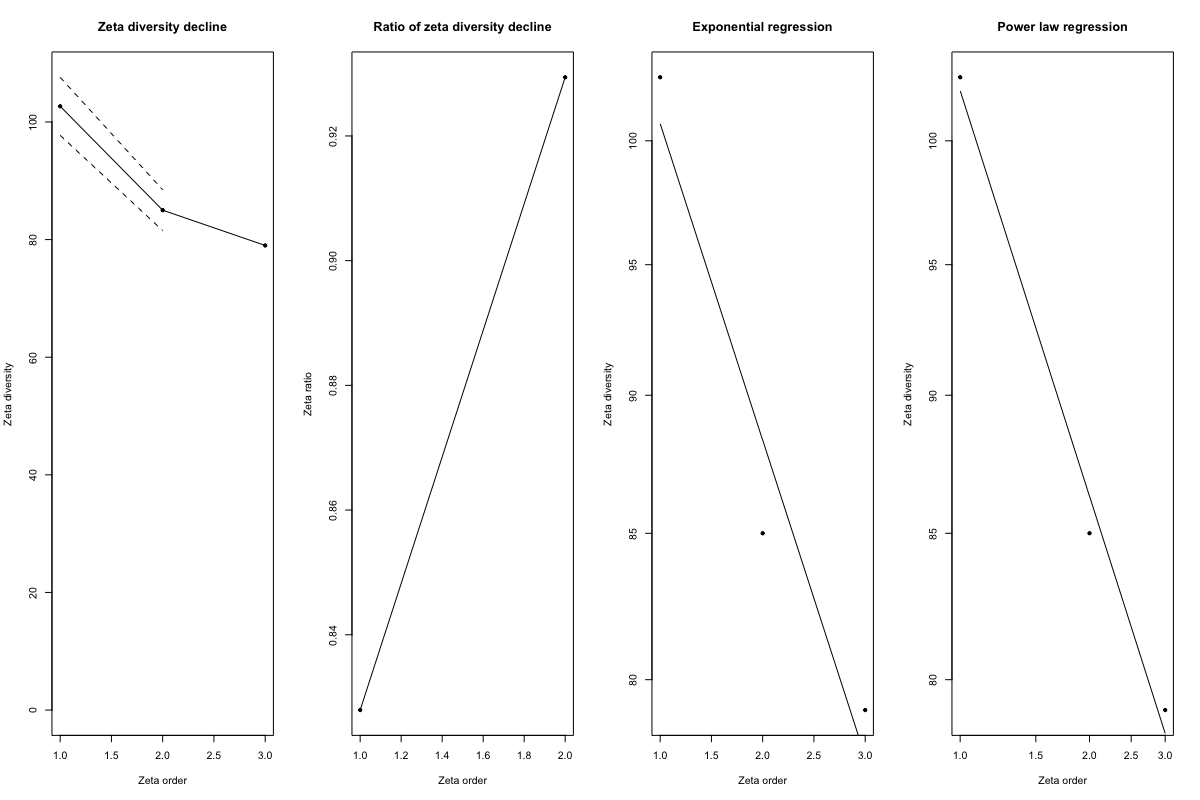

**Figure S20. Zeta Decay of Sites from Bantam-Bintang.**

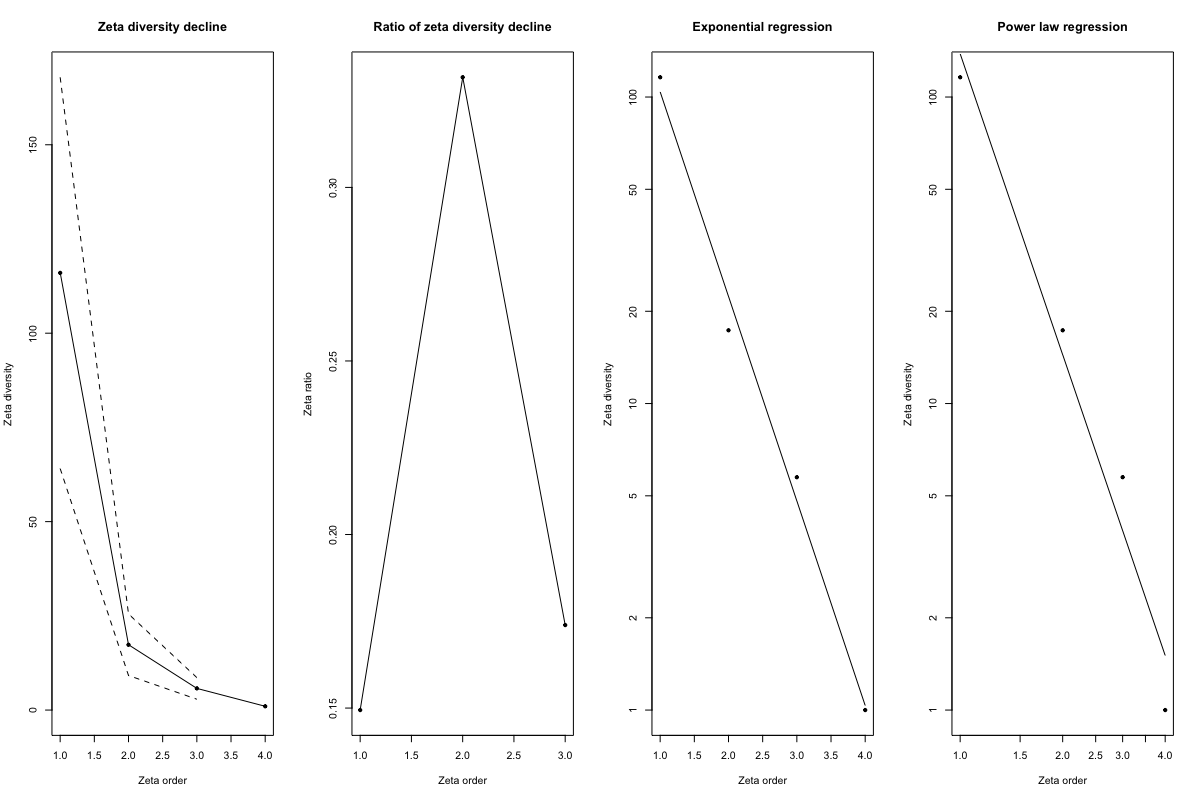

**Figure S21. Zeta Decay of Sites from Derawan.**

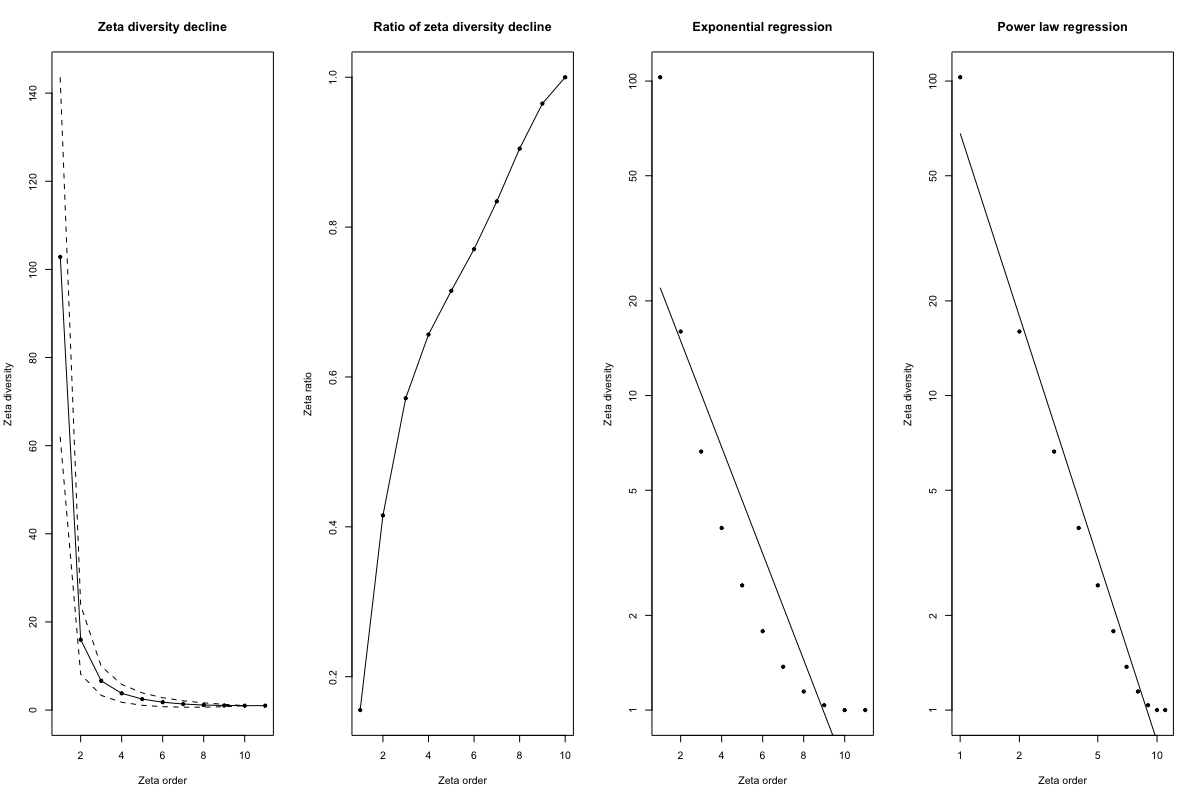

**Figure S22. Zeta Decay of Sites from Lembeh Strait.**

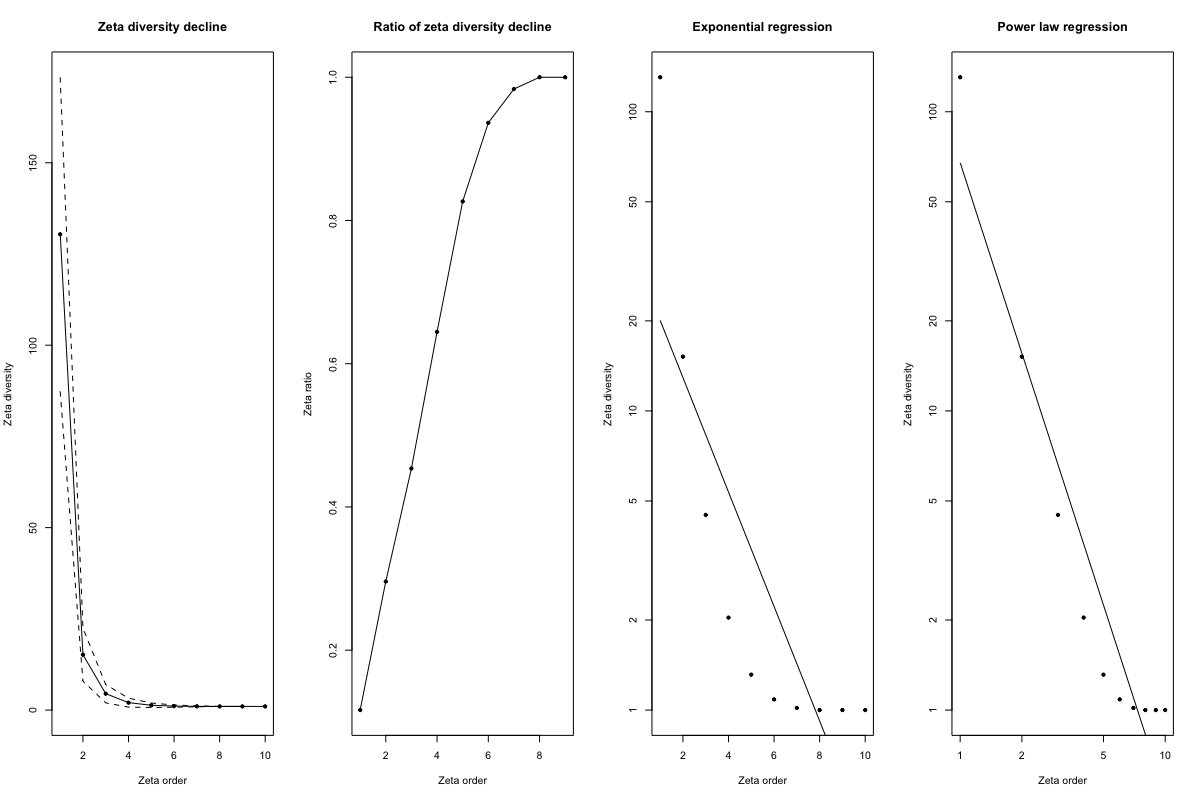

**Figure S23. Zeta Decay of Sites from Raja Ampat.**

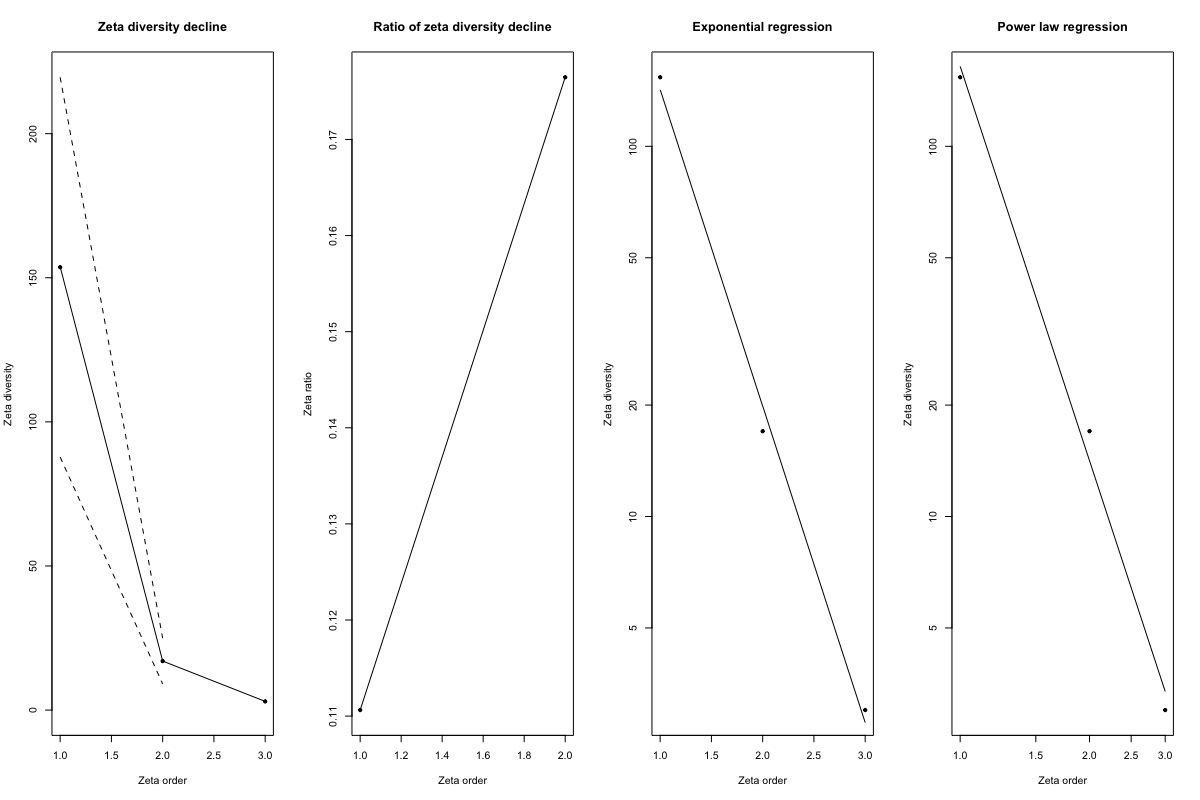

**Figure S24. Zeta Decay of Sites from Ternate.**

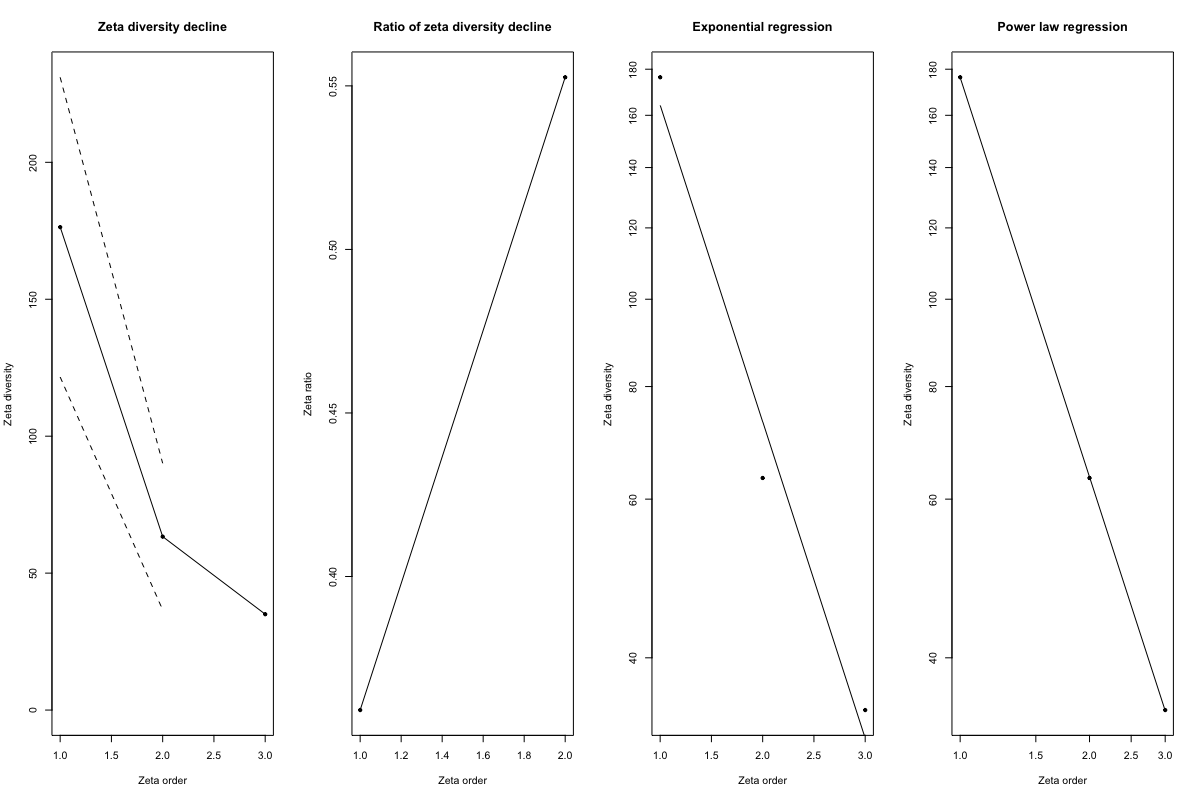

**Figure S25. Zeta Decay of Sites from Wakatobi.**
