## Supplementary material for "Environmental DNA in a Global Biodiversity Hotspot: Lessons from Coral Reef Fish Diversity Across the Indonesian Archipelago": Figures

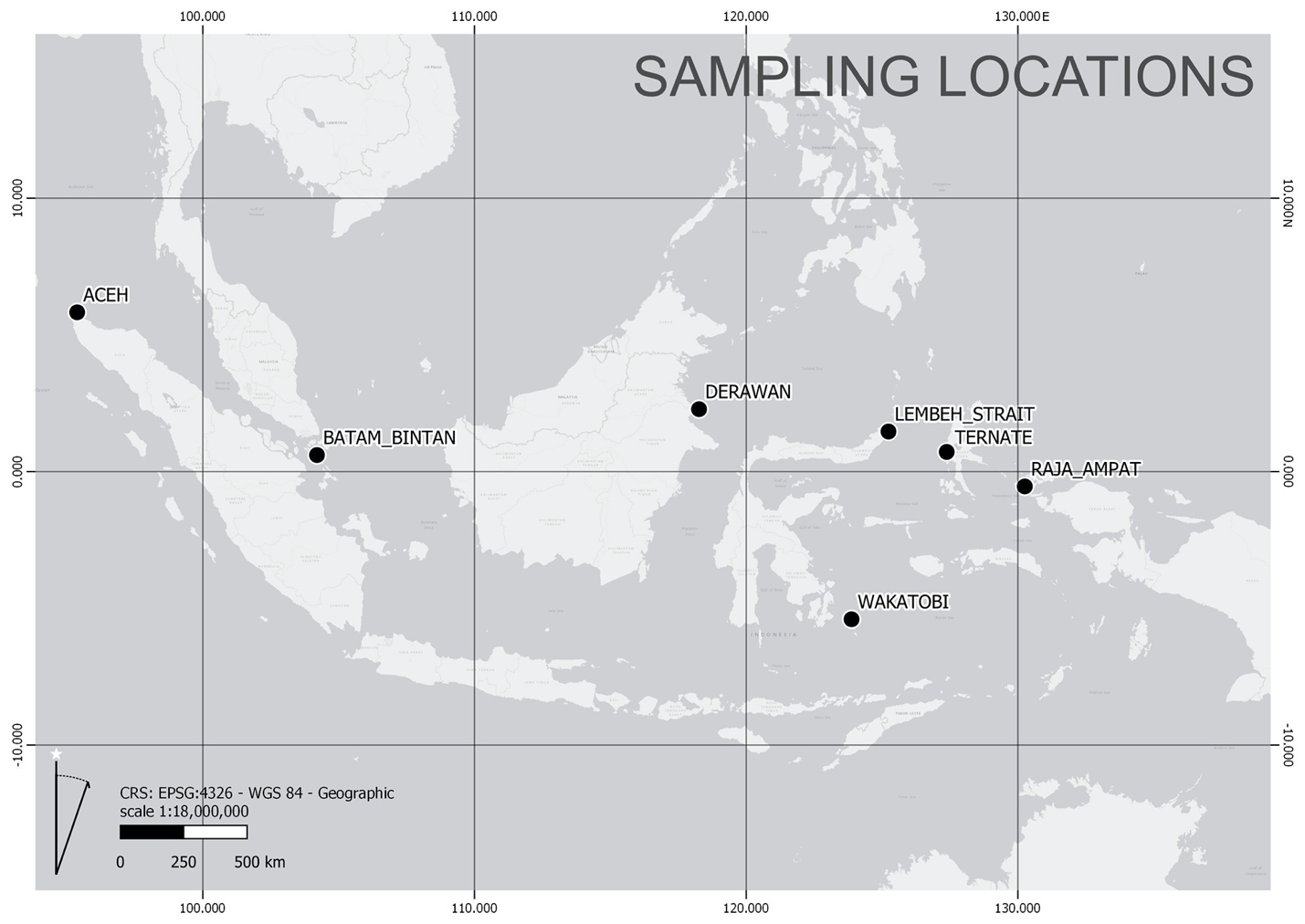

**Figure 1**. eDNA sampling locations across the Indonesian Archipelago.

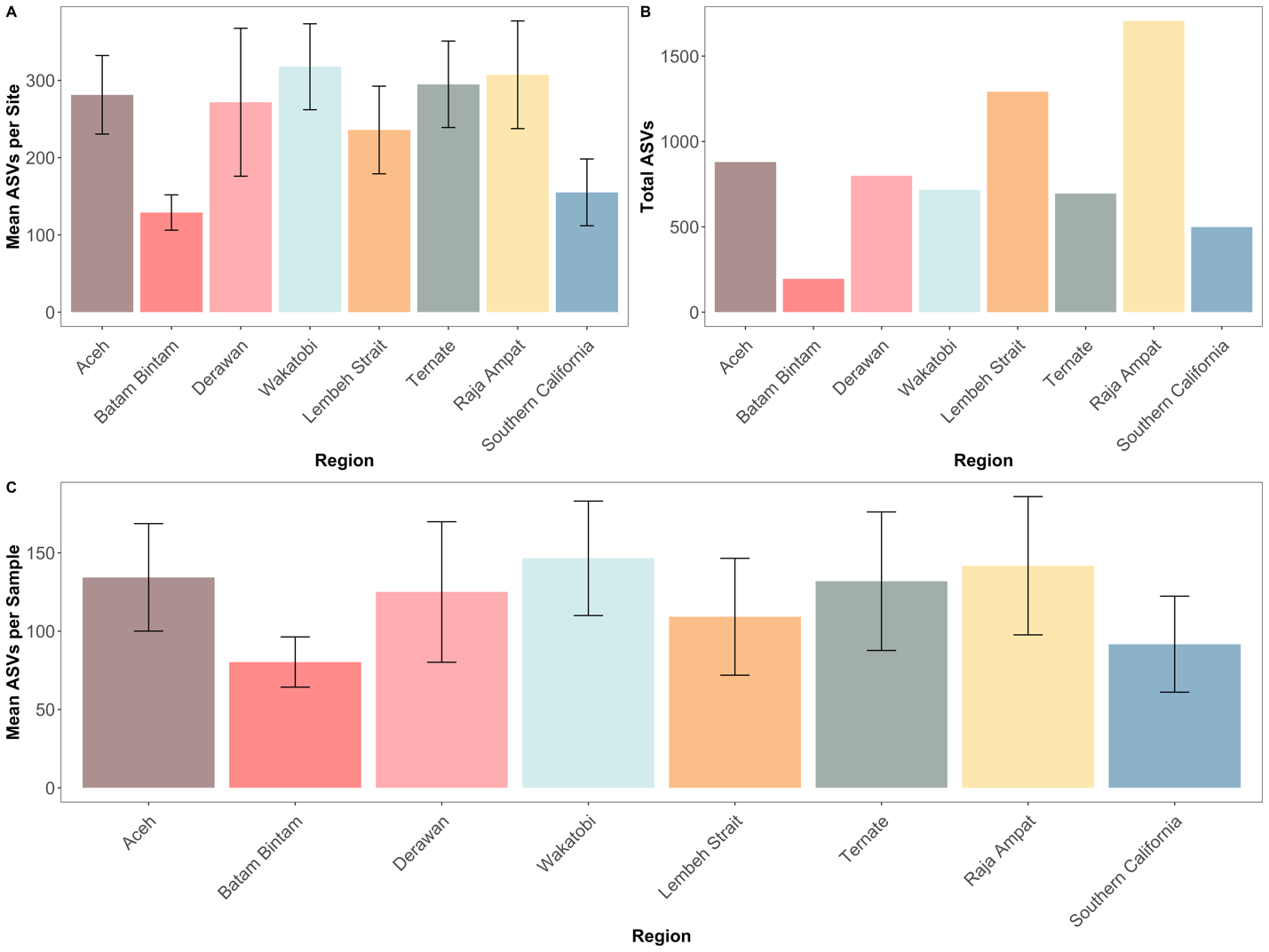

**Figure 2**. Amplicon Sequence Variant (ASV) Richness Per Sample, Site, and Region.

Mean Amplicon Sequence Variant (ASVs) richness per Site (Panel A), Total ASV richness (Panel B), and mean ASV richness per one liter sample replicates (Panel C). Although Indonesia has substantially higher fish diversity than Southern California at the region and site level, mean ASVs per one liter sample are similar between samples collected across Indonesia and Southern California. Colors depict different regions sampled.

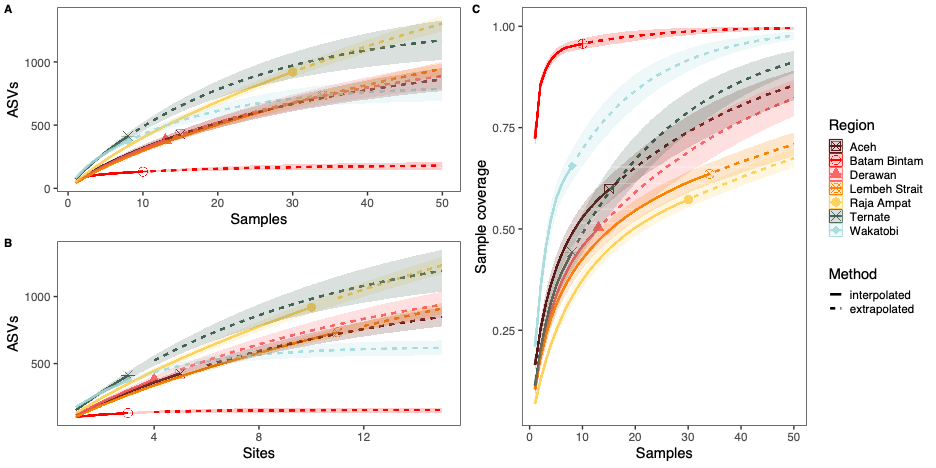

**Figure 3.** Amplicon Sequence Variances (ASVs) Accumulation Curves.

Amplicon Sequence Variants (ASVs) accumulation curves across one liter sample replicates (Panel A) and within each site (Panel B) as well as sample coverage estimates (Panel C)

demonstrate under sampling of Indonesia marine fish diversity through eDNA metabarcoding. Accumulation curves were generated using the iNext package in R. Colors depict different regions samples and line weight depicts interpolated or extrapolated data.

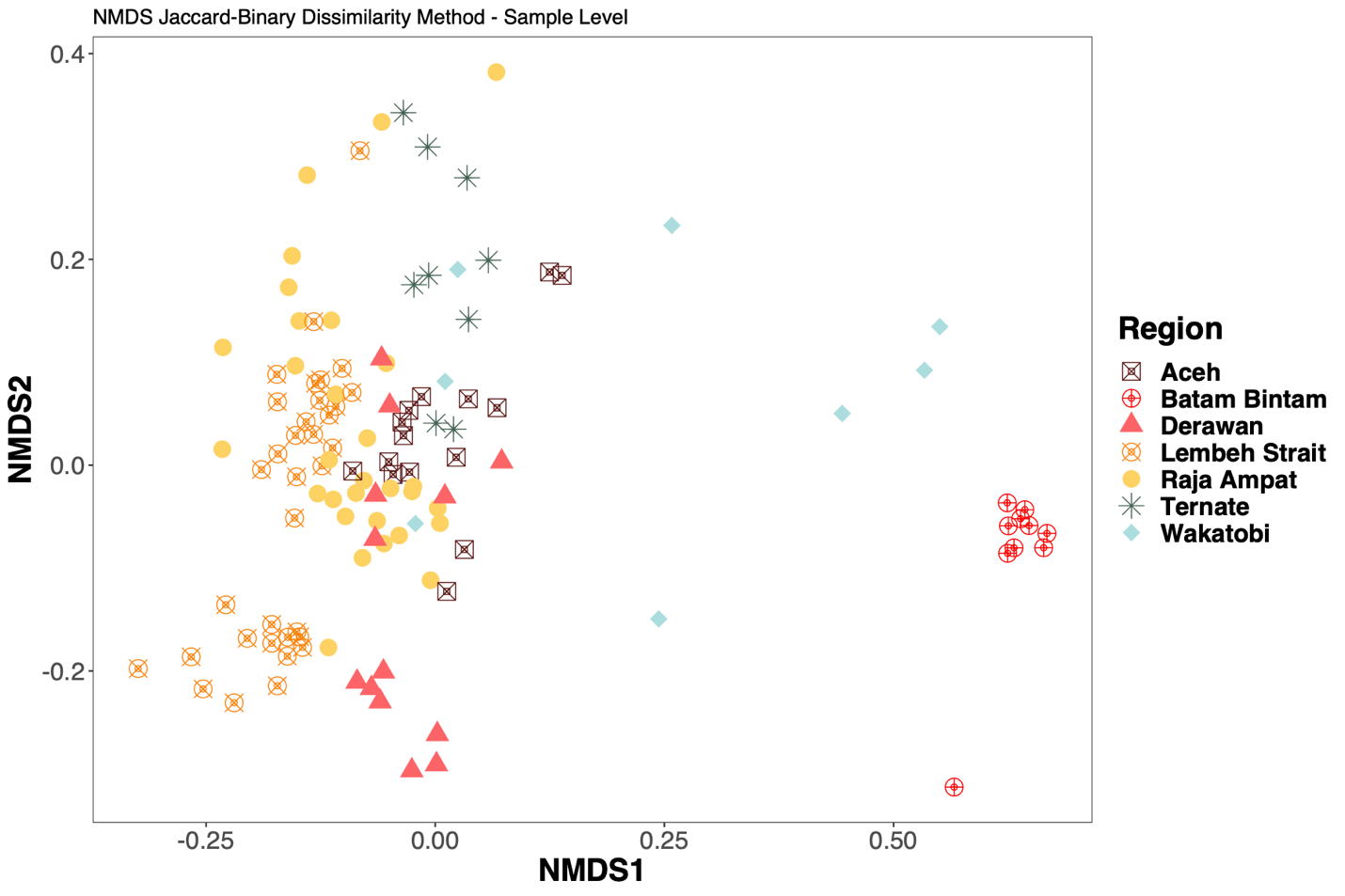

**Figure 4. Nonmetric Multidimensional Scaling (NMDS) Ordination.**

Ordination of Nonmetric Multidimensional Scaling (NMDS) visualizing jaccard-binary fish community similarities between sites and regions. Ordination was implemented through the phyloseq and vegan packages in R. Shapes and colors depict regions sampled.

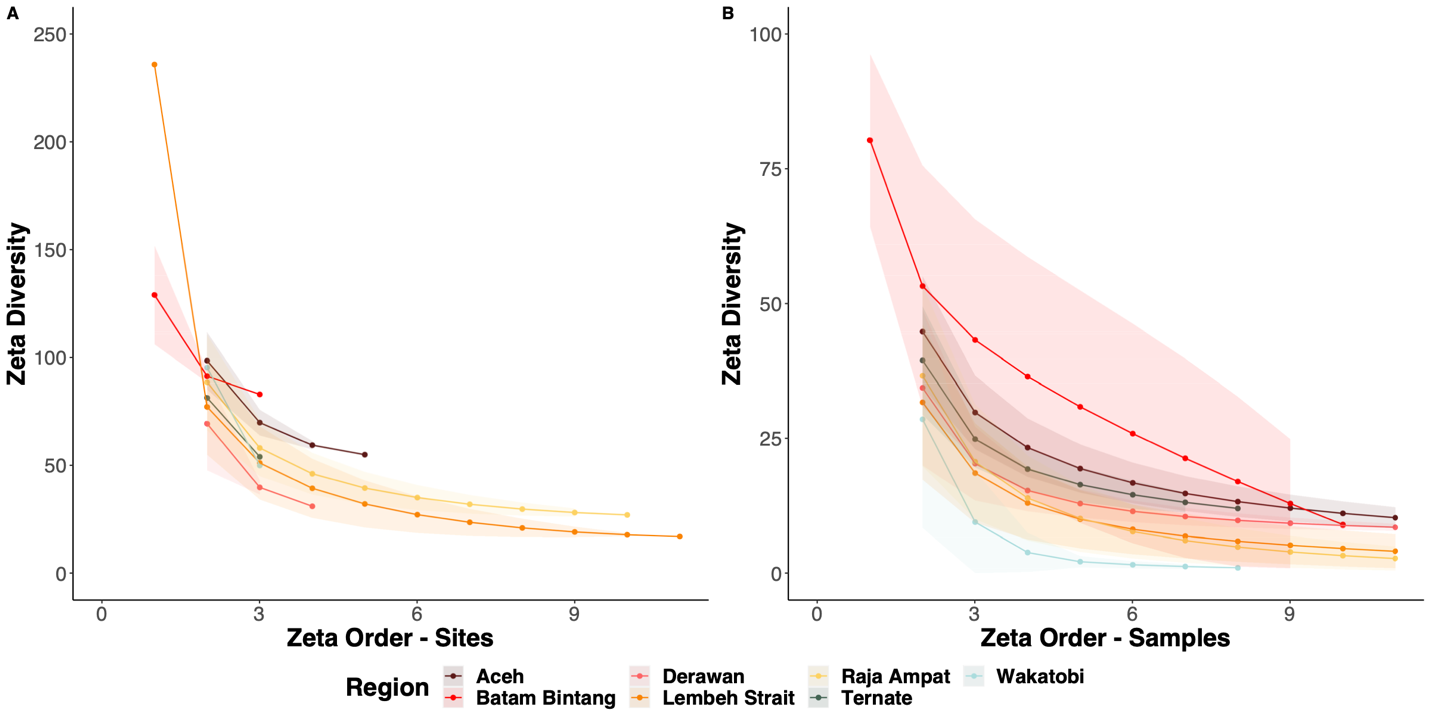

**Figure 5. Zeta Diversity Decay Across Sites and Samples.**

Zeta diversity decay of ASVs across one liter sample replicates and sites. Western regions (Aceh and Batam Bintang) show slower decay than Eastern regions in the Coral Triangle indicating lower community turnover across sites (Panel A) and sample replicates (Panel B).

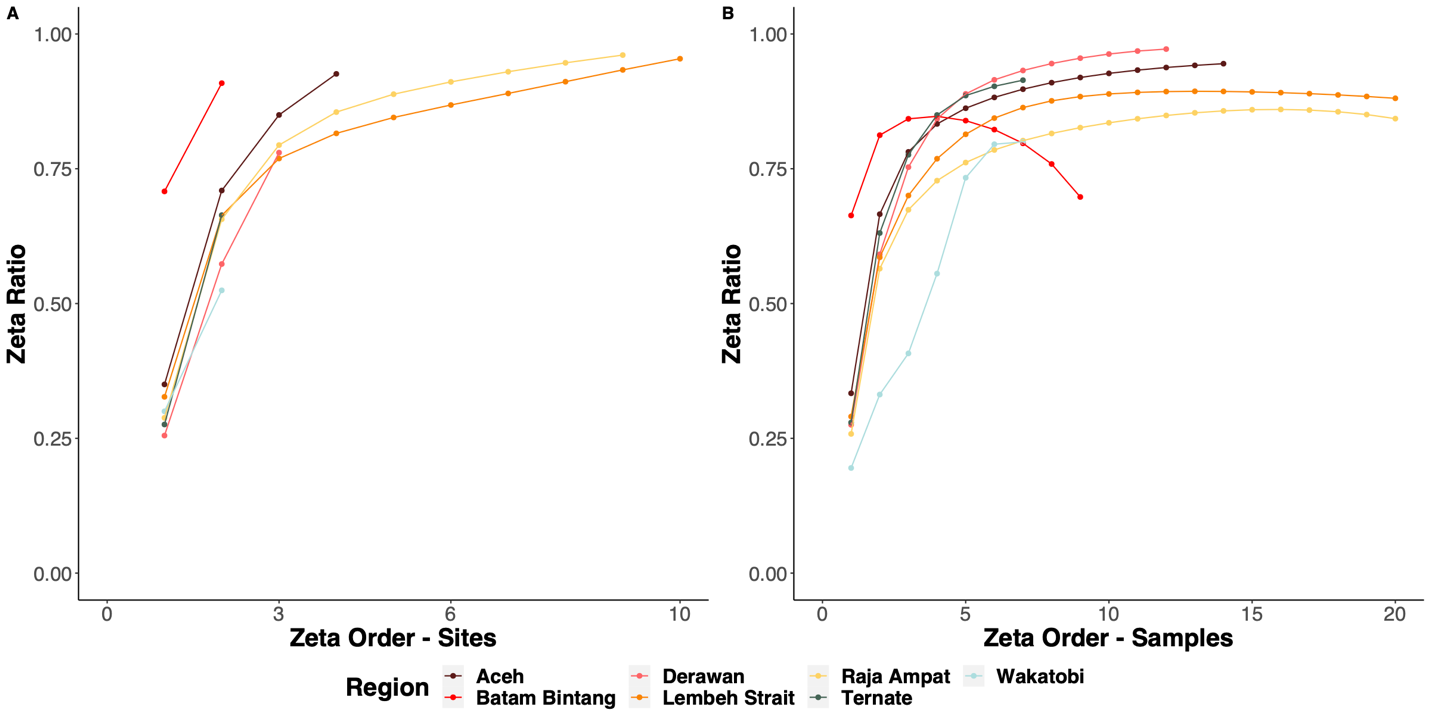

**Figure 6. Species Retention Rates Across Sites and Samples.**

Species retention rates (zeta ratio of ASVs) across one liter sample replicates and sites. Sites in the two Western regions (Aceh and Batam Bintang) show higher ASV retention than Eastern regions in the Coral Triangle indicating lower community turnover across sites (Panel A). Eastern sites in the heart of the Coral Triangle, particularly Raja Ampat and Lembeh Strait, have lower ASV retention rates across samples than most other sites across all zeta orders indicating fewer common species shared across increasing sample replication (Panel B). The bell shaped curve of Batam Bintang is indicative of a few common species shared across samples followed by community turnover, although at the site level there is high ASV retention.
