## Supplementary material for "Environmental DNA in a Global Biodiversity Hotspot: Lessons from Coral Reef Fish Diversity Across the Indonesian Archipelago": Tables

Table 1. Number of sequences reads per region and location. R1 is summary sequence reads produced by Anacapa pipeline, and R2 is the number of high-quality sequences reads (after removing blanks, negative and positive controls, non-marine fishes, singletons, and after standardizing sequence number across all samples)

| **Region** | **Data Processing Step** | **Number of Sites** | **Number of Samples** | **Number of reads** | **Number of ASVs** | **Percent of ASVs identified to species (%)** |
| --- | --- | --- | --- | --- | --- | --- |
| Aceh | R_1_ | 5 | 15 | 1,957,985 | 5421 | 68.4% |
|  | R_2_ | 5 | 15 | 1,719,514 | 879 | 61.9% |
| Batam-Bintan | R_1_ | 3 | 10 | 1,504,845 | 3939 | 84.3% |
|  | R_2_ | 3 | 10 | 1,236,154 | 196 | 85.7% |
| Derawan | R_1_ | 4 |  | 1,259,394 | 3897 | 65.9% |
|  | R_2_ | 4 | 13 | 1,058,236 | 799 | 57.7% |
| Wakatobi | R_1_ | 3 | 8 | 1,153,275 | 6110 | 84.0% |
|  | R_2_ | 3 | 8 | 1,095,430 | 717 | 65.7% |
| Lembeh Strait | R_1_ | 11 | 34 | 3,197,936 | 9312 | 64.6% |
|  | R_2_ | 11 | 34 | 2,904,257 | 1292 | 65.6% |
| Ternate | R_1_ | 3 | 9 | 1,385,498 | 4786 | 76.5% |
|  | R_2_ | 3 | 9 | 1,301,880 | 695 | 66.2% |
| Raja Ampat | R_1_ | 10 | 30 | 4,030,226 | 14898 | 69.8% |
|  | R_2_ | 10 | 30 | 3,624,219 | 1706 | 62.9% |

Table 2. Comparison of fish diversity in Raja Ampat based on 1) total ASVs, 2) total Species from eDNA, 3) total Species in Visual Census, and 4) total Species Overlap

| **Region** | **Site** | **Total ASVs** | **Species from eDNA** | **Species in Visual Census** | **Species Overlap** | **Percent Overlap** |
| --- | --- | --- | --- | --- | --- | --- |
| Raja Ampat | Anita Garden | 324 | 60 | 218 | 25 | 11.0% |
|  | Cape Kri | 241 | 36 | 272 | 17 | 6.2% |
|  | Kri Lagoon | 315 | 229 | 299 | 51 | 12.0% |
|  | Kabui Strait | 833 | 112 | 81 | 22 | 14.7% |
|  | Melissa Garden | 272 | 84 | 214 | 34 | 14.7% |
|  | Secret Lagoon | 260 | 75 | 89 | 11 | 7.7% |
|  | Sardine Reef | 323 | 77 | 210 | 37 | 17.4% |
|  | West Mansuar | 184 | 40 | 227 | 16 | 6.8% |
